# Analysing the safe and just operating space of agriculture in the world: past, present and future

**DOI:** 10.1101/824797

**Authors:** Ajishnu Roy, Kousik Pramanick

## Abstract

Agriculture, along with industry and household sector are three major sectors of human consumption. Agriculture has proved to be a major contributor to exceeding planetary boundaries. Here, we have explored the impact of agriculture in the Earth system processes, through eight dimensions of planetary boundaries or safe operating spaces: climate change (10.73%), freshwater use (91.56%), arable land use (37.27%), nitrogen use (95.77%), phosphorus use (87.28%), ecological footprint (19.42%), atmospheric pollution (2.52% - 38.08%) and novel entities. In this work, we have also shown role of agriculture to the socio-economic development dimensions: gender equality, employment and economic growth. We have shown that the safe operating limits for agriculture are going to decline by almost 55% (climate change), 300% (freshwater use), 50-55% (arable land use), 180% (nitrogen use), 265% (phosphorus use) and 20% (ecological footprint) in 2050, if the most inefficient way of consumption is chosen and continued. To alleviate the role of agriculture in transgressing planetary boundaries, it is indispensable to comprehend how many roles of agriculture is playing and where which target should be set to framework the national agricultural policies in coherence with attaining sustainable development goals of UN by 2030.

## 1. Introduction

The World has now arrived the Anthropocene era in which human actions, above all others, have the most profound impact on Earth system functioning (Dirzo et al., 2014, Steffen et al., 2018). The significant influence of agriculture on the environment is well acknowledged (Tilman 1999, Tilman et al. 2001, Foley et al. 2005). Agriculture interacts with global earth system processes through numerous pathways (viz. greenhouse gas emissions, excessive water use, land-use change, and biodiversity impacts). The global agriculture is a chief driver of climate change (Vermeulen et al., 2012, Wollenberg et al., 2016, Carlson et al., 2017, Campbell et al., 2018), depletion of freshwater resources (Destouni et al., 2013), land-use system change (Zabel et al., 2014, Scown et al., 2019, Stehfest et al., 2019), biogeochemical flows of nitrogen (Galloway et al., 2008, Liu et al., 2010, Robertson et al., 2010, De Vries et al., 2013) and phosphorus (Cordell et al., 2014, Zhang et al, 2017) (through fertilizer and manure application), biodiversity loss (Newbold et al., 2016, Mace et al., 2018), emission of atmospheric pollutants and introduction of novel entities (Stehle et al., 2015). Agriculture has contributed to the exceeding of several of the proposed ‘planetary boundaries’ which define a safe operating space for humankind on a stable Earth system (Campbell et al., 2017, Conijn et al., 2018, Springmann et al., 2018). If this current agricultural consumption pattern continues, the biophysical pressures of the agriculture would deepen, and humanity might come near and cross the planetary boundaries. Beyond those safe limits, key ecosystem functions could be destabilized on which, not only billions of human populations, but also all the organisms that comprise the biodiversity, depend on (Rockström et al., 2009, Steffen et al., 2015, Scheffer et al., 2015, O’Neill et al., 2018).

However, there are a few points that need to be recognized which the purpose of this work was. They are - (1) categorization of the available safe limits (per capita scale) of planetary boundaries to be integrated into the sector of agriculture, (2) appropriating the trends of these parameters of biophysical consumption of agriculture-related to PB, based on availability of data, (3) as per the latest year of available data, representation of countries which are either exceeded or under respective safe limits of each dimension of PB, (4) the input of agriculture (global to national level) in each dimension of PB framework, (5) appropriation of amount of biophysical resources that are going to be consumed in agriculture according to different scenarios up to 2050 and their comparison, (6) based on population projection of different scenarios available, how the per capita safe limits of agricultural biophysical consumption is going to change up to 2050, (7) comprehension of the trends of indicators related to socioeconomy of agriculture and their relation to biophysical consumption dimensions.

## 2. Data and Method

### 2.1 Planetary boundaries

#### 2.1.1 Climate change

As per Emissions Gap Report (UNEP, 2018), emissions of all greenhouse gases are not to exceed 24 GtCO_2_-e (range 22-30 GtCO_2_-e) in 2030 if the <1.5 □ target is to be attained in 2100 with >66% chance (Table 3.1, page 19). Diving this 24 GtCO_2_-e with the global population we get per capita global scale Climate change boundary of 3.29 (3.01-4.11) tCO2-e year^-1^ (for 2014). This could be used as safe climate change limit for all sectors, not specific to agriculture. But according to van Vuuren et al. (2011), Wollenberg et al. (2016) and Springmann et al. (2018), food-related GHG emission budget is 4.7 (4.3-5.3) GtCO_2_-e, which can be calculated to per capita global sector-specific boundary of 0.64 (0.59-0.73) tCO_2_-e year^-1^ (for 2014). We have calculated the stake of agricultural GHG emission in total, to get the contribution of agriculture in climate change PB.

#### 2.1.2 Freshwater use

Gerten et al. (2013) have updated freshwater use PB to 2800 km^3^ y^-1^(with avg. uncertainty range: 1100-4500 km^3^ y^-1^). We have divided 2800 km^3^ water with the global population to get the global average per capita scale freshwater use boundary of 370.85 (145.69-596) m^3^ y^-1^ (2017). This could be used as safe freshwater use limit for all sectors, including agriculture. However, based on works of Shiklomanov & Rodda (2004) and Springmann et al. (2018), food-related freshwater use (Blue water) budget is 1980 (780-3190) km^3^, which can be calculated to 262.24 (103.31-422.5) m^3^ y^-1^ (2017). We have calculated the share of agricultural freshwater use in total, from the Aquastat database, to get the contribution of agriculture in freshwater use PB. We have also used another 3 indicators related to water footprint (WF). These are – (1) agricultural WF of consumption (% of total), (2) agricultural blue WF of consumption (% of total blue WF) and (3) agricultural grey WF of consumption (% of total grey WF).

#### 2.1.3 Arable land use

The planetary boundary of land use is <15% of global ice-free land cover converted per year to cropland (i.e. 1995 Mha) (Rockström et al., 2009). We have divided this 1995 Mha with the global population to get a global per capita scale average land use PB of 0.27 ha year^-1^ (2015). This could be used as safe land use limit for all sectors. According to Springmann et al. (2018), food-related cropland use budget is 12.6 (10.6-14.6) Mkm^2^, which can be calculated to 0.17 (0.14-0.2) ha capita^-1^ year^-1^ (2015). We have calculated the share of agricultural land use in total, to get the contribution of agriculture in land use PB.

#### 2.1.4 Nitrogen use

According to Steffen et al. (2015), the planetary boundary of global N flow is 62 Tg N per year. We have divided 62 Tg N y^-1^ with the global population to get the global average per capita scale boundary of 8.4kg N year^-1^ (2015). This could be used as safe nitrogen use limit for all sectors. Based on works of Mueller et al. (2012), de Vries et al. (2013) and Springmann et al. (2018), food-related nitrogen use budget is 69 (52-113) TgN, which can be calculated to 9.34 (7.04-15.3) kg N capita^-1^ year^-1^ (2015). We have calculated the share of agricultural nitrogen use in total, to get the contribution of agriculture in nitrogen use PB.

#### 2.1.5 Phosphorus use

According to Steffen et al. (2015), the planetary boundary of global phosphorus flow (mined and applied to erodible or agricultural soils) is 6.2 Tg P y^-1^. We have divided 6.2 Tg P y^-1^ with the global population to get the global average per capita scale boundary of 0.84kg P year^-1^ (2015). This could be used as safe phosphorus use limit for all sectors. But, based on works of Mueller et al. (2012) and Springmann et al. (2018), food-related phosphorus use budget is 16 (8-17) Tg P, which can be calculated to 2.17 (1.08-2.3) kg P capita^-1^ year^-1^(2015). We have calculated the share of agricultural phosphorus use in total, to get the contribution of agriculture in phosphorus use PB.

#### 2.1.6 Ecological footprint (EF)

According to the Global Footprint Network (GFN), the world has 12 billion ha biologically productive land and sea area. We have divided 12 billion ha with the global population to get the global average per capita scale boundary of 1.6gha year^-1^ (2016). We have calculated the share of agricultural ecological footprint (i.e. cropland EF component) in total, to get the contribution of agriculture in ecological footprint PB.

#### 2.1.7 Air pollution

We have used 10 different indicators to get the overall view about the contribution of agriculture to air pollution. These are – ammonia (NH3), black carbon (BC), carbon monoxide (CO), nitrogen oxides (NOx), non-methane volatile organic compounds (NMVOC), organic carbon (OC), particulate matter 2.5 (PM _2_._5_) – bio and fossil origin, particulate matter 10 (PM _10_) and sulphur dioxide (SO_2_). We have calculated the share of agricultural air pollutant emission in total, to get the contribution of agriculture in atmospheric pollution PB.

#### 2.1.8 Novel entities

To appropriate this dimension, we have used 3 indicators – pesticide use in cropland (kg/ha), pesticide total use (active ingredients) (kg/capita) and synthetic fertilizer use (kg/capita). As the boundaries are yet to be set, we have only been able to use the per capita values.

**Table 1:**
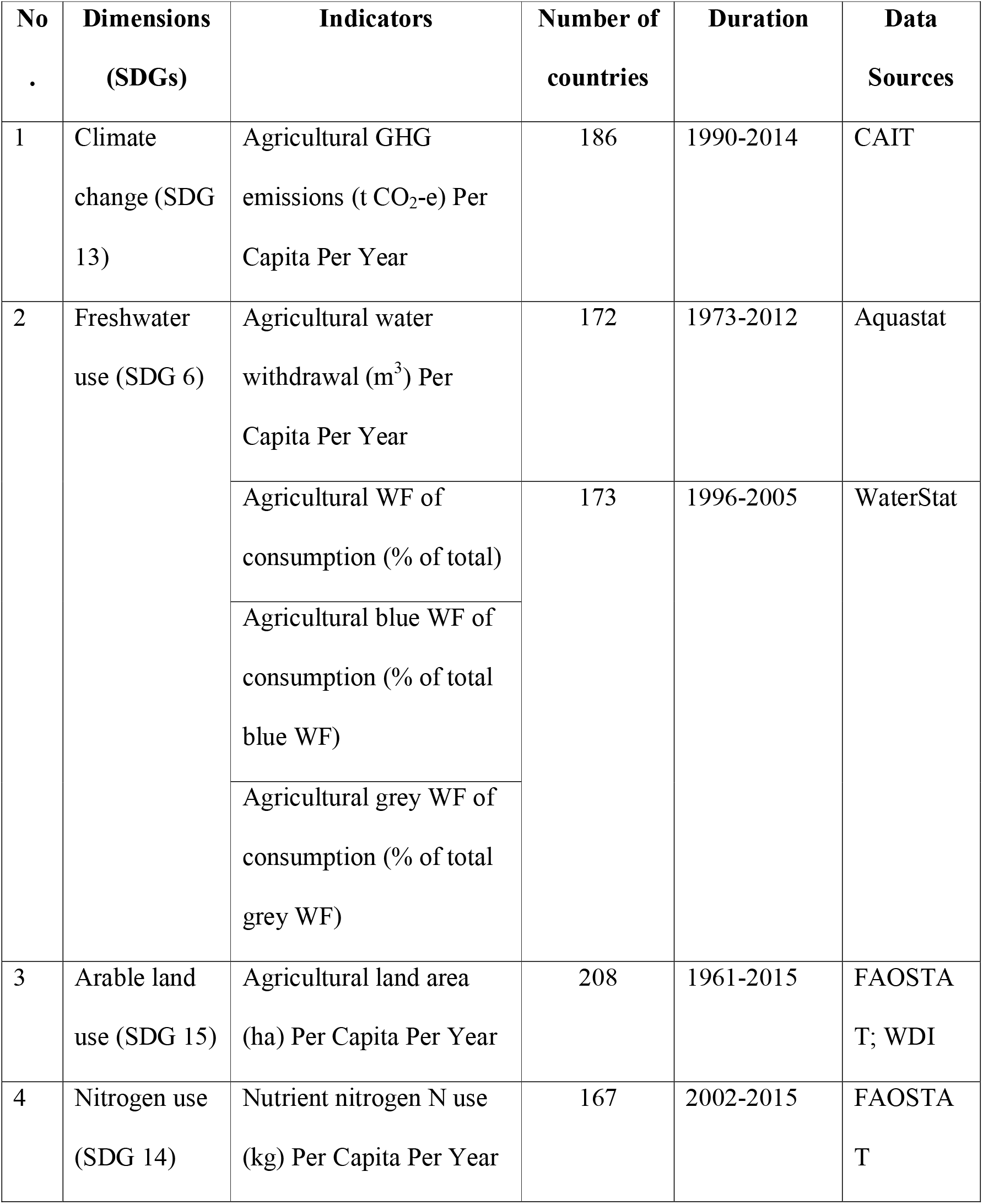
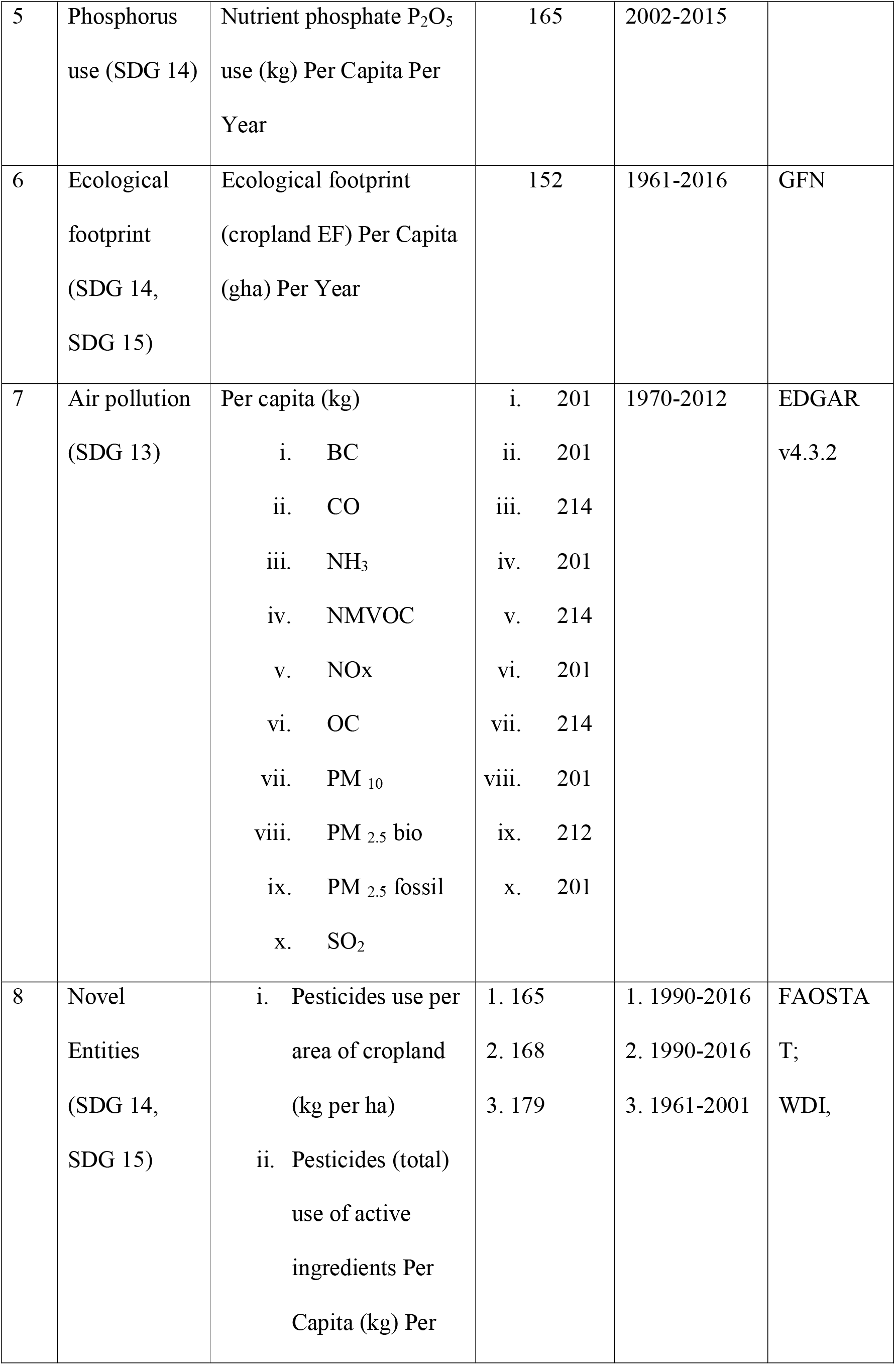
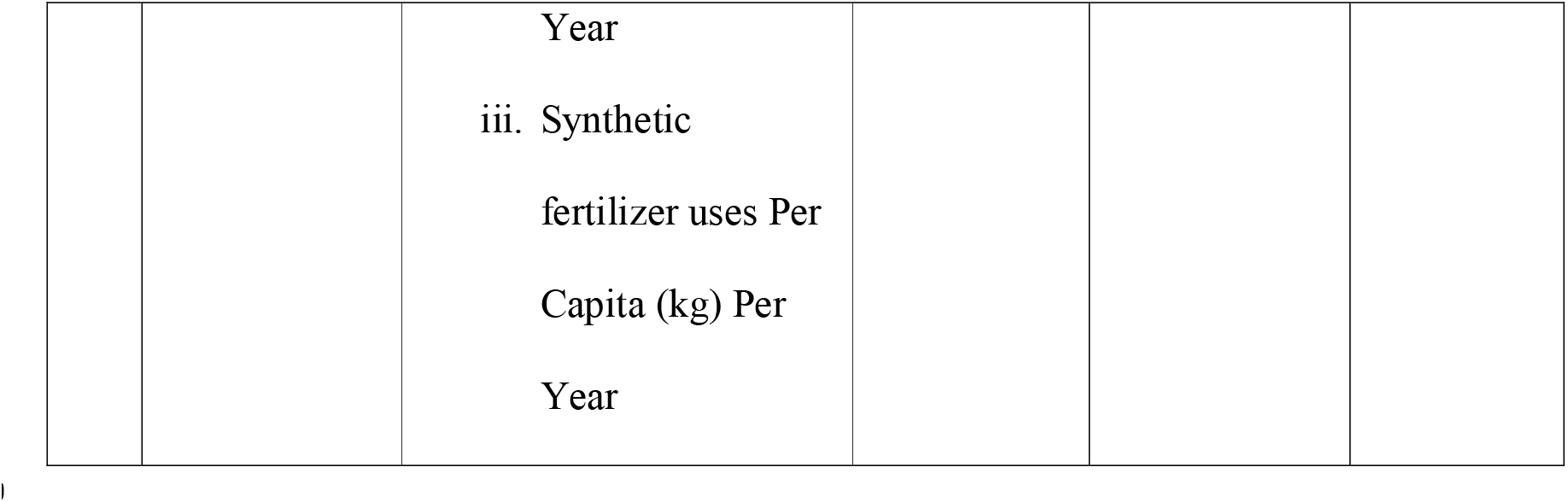
Biophysical indicators of planetary boundaries related to agriculture.

| No<br>. | Dimensions<br>(SDGs) | Indicators | Number of<br>countries | Duration | Data<br>Sources |
| --- | --- | --- | --- | --- | --- |
| 1 | Climate<br>change (SDG<br>13) | Agricultural GHG<br>emissions (t CO<sub>2</sub>-e) Per<br>Capita Per Year | 186 | 1990-2014 | CAIT |
| 2 | Freshwater<br>use (SDG 6) | Agricultural water<br>withdrawal (m<sup>3</sup>) Per<br>Capita Per Year | 172 | 1973-2012 | Aquastat |
|  |  | Agricultural WF of<br>consumption (% of total) | 173 | 1996-2005 | WaterStat |
|  |  | Agricultural blue WF of<br>consumption (% of total<br>blue WF) |  |  |  |
|  |  | Agricultural grey WF of<br>consumption (% of total<br>grey WF) |  |  |  |
| 3 | Arable land<br>use (SDG 15) | Agricultural land area<br>(ha) Per Capita Per Year | 208 | 1961-2015 | FAOSTA<br>T; WDI |
| 4 | Nitrogen use<br>(SDG 14) | Nutrient nitrogen N use<br>(kg) Per Capita Per Year | 167 | 2002-2015 | FAOSTA<br>T |
| 5 | Phosphorus use (SDG 14) | Nutrient phosphate P<sub>2</sub>O<sub>5</sub> use (kg) Per Capita Per Year | 165 | 2002-2015 |  |
| 6 | Ecological footprint (SDG 14, SDG 15) | Ecological footprint (cropland EF) Per Capita (gha) Per Year | 152 | 1961-2016 | GFN |
| 7 | Air pollution (SDG 13) | Per capita (kg)<br>i. BC<br>ii. CO<br>iii. NH<sub>3</sub><br>iv. NMVOC<br>v. NO<sub>x</sub><br>vi. OC<br>vii. PM<sub>10</sub><br>viii. PM<sub>2.5</sub> bio<br>ix. PM<sub>2.5</sub> fossil<br>x. SO<sub>2</sub> | i. 201<br>ii. 201<br>iii. 214<br>iv. 201<br>v. 214<br>vi. 201<br>vii. 214<br>viii. 201<br>ix. 212<br>x. 201 | 1970-2012 | EDGAR v4.3.2 |
| 8 | Novel Entities (SDG 14, SDG 15) | i. Pesticides use per area of cropland (kg per ha)<br>ii. Pesticides (total) use of active ingredients Per Capita (kg) Per | 1. 165<br>2. 168<br>3. 179 | 1. 1990-2016<br>2. 1990-2016<br>3. 1961-2001 | FAOSTA T; WDI, |
|  |  | Year |  |  |  |
|  |  | iii. Synthetic |  |  |  |
|  |  | fertilizer uses Per |  |  |  |
|  |  | Capita (kg) Per |  |  |  |
|  |  | Year |  |  |  |

### 2.2 Future scenario

As we have calculated all of the biophysical indicators on per capita basis, it is conceivable to project probable future scenario of total biophysical consumption from agricultural sector. We have collected future population projection (2015-2050) data (median range prediction value of 50%) of the World from the WUP, UN (2018 Revision) and also projections according to five shared socioeconomic pathways (SSPs); incorporating these with per capita consumption indicators (n = 9) of planetary boundaries. We have calculated 12-18 projection series for each dimension of PB, (i) with the lowest value (which has happened in the past year), (ii) highest value (which has happened in the past year) and (iii) business-as-usual (BAU) scenario with the latest year of available data.

### 2.3 Doughnut economy of agriculture

We have also analysed various socioeconomic factors related to the agriculture, as – income (via GDP and GNI per capita), employment, value added by agriculture and share of agriculture in male and female employment, which are in turn connected to UN SDG 1, 5 and 8; and 3 dimensions of doughnut economy (income, job, gender equality) (Raworth, 2017).

**Table 2:**
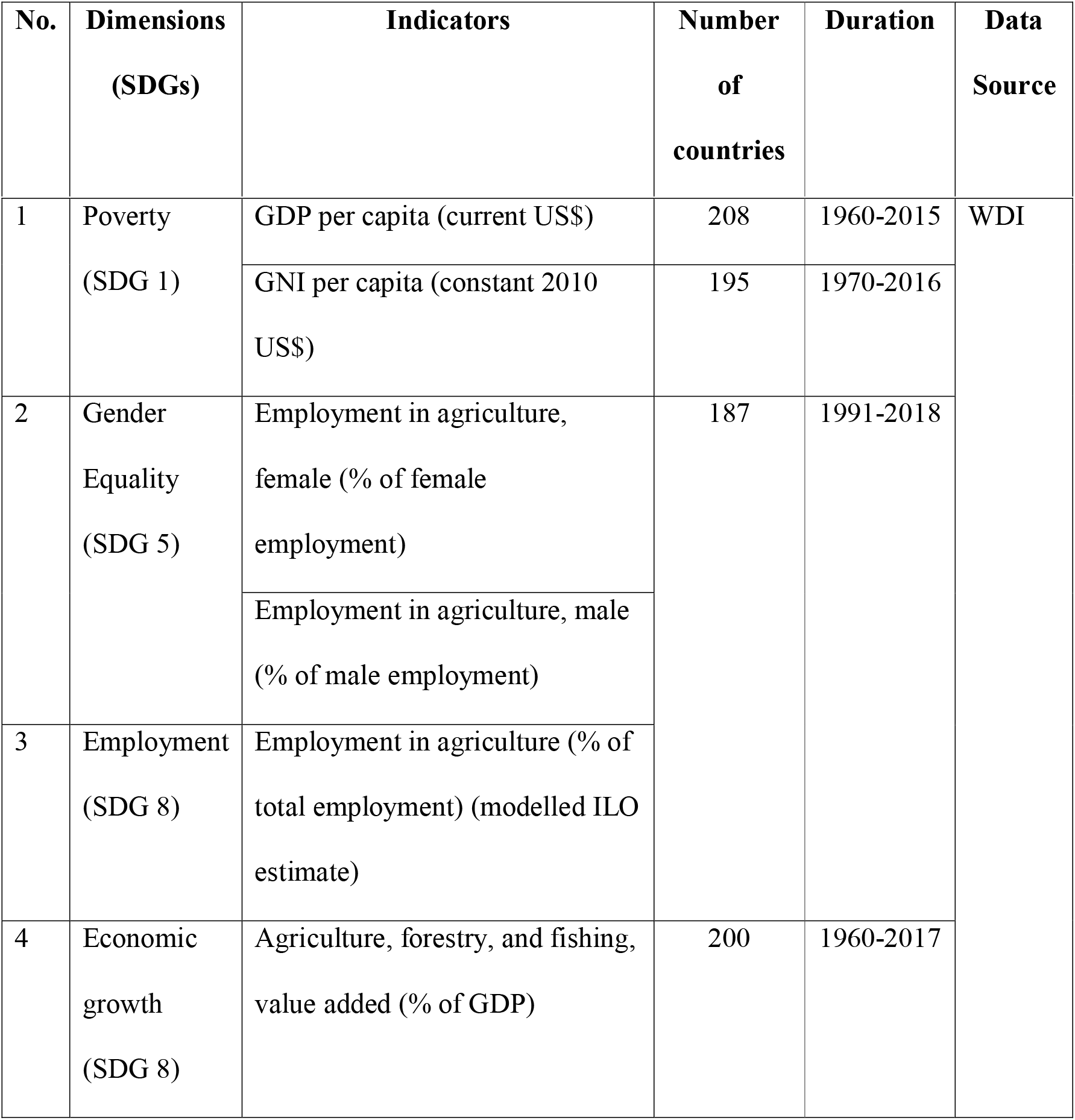
Socioeconomic development indicators related to agriculture.

## 3. Results

### 3.1 Planetary boundaries

#### 3.1.1 Climate change

Agricultural GHG emission has decreased in the world (16.59%), Africa (23.99%), Asia (5.87%), Europe (36.66%), North America (15.61%) and Oceania (37.03%) from 1990 level. This agricultural GHG emission is lower than global average per capita climate change PB in Alliance of Small Island States (5.63%), Asia (16.34%) and Middle East and North Africa (42.06%); but higher in the world (12.76%), Africa (7.74%), Europe (24.35%), North America (82.35%), Oceania (673.14%). Agricultural GHG emission in other nation groups are also higher than climate change PB (viz. EU – 30.27%; G7 – 33.01%; G77 and China – 5.12%; Latin America and the Caribbean – 128.82%, LDCs – 29.5%; sub-Saharan Africa – 16.04%; UNFCC Annex I – 47.22%; UNFCC non-Annex I – 4.96%) (Supplementary fig 1). The contribution of agriculture to total GHG emission has decreased in the world (2.8%) but increased in the EU (0.84%) since 1990. Agricultural GHG emission used to be high in countries of South Asia, Africa and South American countries in 1990. Though the contribution of agriculture has reduced over time, those same countries still high than in other countries in the world.

#### 3.1.2 Freshwater use

Agricultural use of freshwater was already higher during 1970-1975 in various parts of the world, like – central and South America, most of Africa, all parts of Asia (except – northern Asia) and Oceania. This value even increased further in all of those areas, only slightly decreased in parts of central Asia & Africa. In total WF of consumption, agriculture’s contribution is now 91.5%, i.e., it has increased by 5.66%, globally. As per the latest available data, agriculture is responsible for 91.56% of blue WF and 51.6% of grey WF of consumption at a global level. In the country level, the lowest value of the contribution of agriculture in WF of consumption has risen from 50% to 61%.

#### 3.1.3 Arable land use

Arable land area (ha per capita) have decreased significantly for the world (53.79%), Africa (62.19%), North America (48.31%), South America (11.62%), Asia (54.16%), Europe (38.8%) and Oceania (42.18%), since the 1961 level. Arable land area (per capita) is lower than land-use PB in Asia (34.04%), but higher in the world (13.61%), Africa (15.82%), North (223.64%) and South America (101.03%), Europe (119.54%) and Oceania (603.19%). Arable land area (ha per capita) has also reduced in EU (33.62%), LDC (60.29%), LLDC (53.02%), SIDS (37.95%), LIFDC (63.44%) and NFIDC (62.28%), in comparison with 1961. Among these groups, SIDS (40.86%), LIFDC (12.37%) and NFIDC (4%) have a lower; and EU (24.49%), LDC (8.09%), LLDC (67.04%) have higher arable land area (per capita) than land-use PB (Supplementary fig 2). Agricultural land area (% of total land area) has increased in the world (1.23%), Arab world (4.26%), Caribbean small states (0.71%), East Asia and Pacific (4.3%), Latin America and Caribbean (9.71%), middle east and north Africa (7.56%), PISS (5.3%), small states (0.14%), South Asia (1.55%), sub-Saharan Africa (2.21%); but decreased in central Europe and Baltics (16.89%), EU (11.15%), Europe and Central Asia (1.2%), Micronesia (0.71%), North America (2.65%), OCED members (5.19%) and other small states (0.27%) than 1961 level (Supplementary fig 3). Agriculture’s share of land use used to be high in almost all over the world except for a few countries in 1961. This situation has remained the same at the present time but deteriorated in countries of Asia and Africa in 2015.

#### 3.1.4 Nitrogen use

Nitrogen use (kg) per capita has increased in the world (12.99%), north (8.07%) and South America (50.26%), Asia (17.95%), Europe (9.15%) and Oceania (9.81%); but decreased in Africa (0.95%) than 2002 level. This value has only decreased in EU (3.69%) and SIDS (8.62%); but increased in LDC (23.23%), LLDC (220.46%), LIFDC (26.71%) and NFIDC (3.17%), in comparison with 2002. Agricultural nitrogen use is lower than per capita nitrogen use PB for agriculture in the world (84.15%), Africa (96.75%), north (54.33%) and South America (83.62%), Asia (84.13%), Europe (79.01%) and Oceania (52.24%). It is also showing similar trend of being lower than per capita nitrogen use PB for agriculture in LDC (70.99%), LLDC (55.15%), SIDS (62.53%), LIFDC (6.2%) and NFIDC (37.95%), except – higher in EU (137.89%) (Supplementary fig 4). Nitrogen use in agriculture (%) is presently more than 95% in all of the continents along with the world, except – North America (78.41%). Also, it has been increasing in all, except –Africa and Oceania (0.05% and 0.09% decrease). Similarly, nitrogen use in agriculture (%) is presently more than 95% in all of the national groups (like - LDC, LLDC, SIDS, LIFDC and NFIDC) (Supplementary fig 5). As this data is available only for a few countries at the national scale, it is difficult to understand overall country-level comparative understanding through longer duration.

#### 3.1.5 Phosphorus use

Phosphorus use (kg per capita) has increased in the world (18.48%), Africa (5.16%), North (2.69%) and South America (51.66%) and Asia (31.39%); but decreased in Europe (10.88%) and Oceania (21.07%) from 2002 levels. Similarly, it has also increased in LDC (49.58%), LLDC (124.98%), SIDS (7.16%), LIFDC (34.01%), NFIDC (21.94%); only decreased in EU (27.53%) from 2002 level (Supplementary fig 6). Most of these groups have crossed the phosphorus use PB for agriculture, like – world (199.34%), Asia (204.84%), Europe (130.02%), North (593.2%) and South America (573.05%), Oceania (1614.97%), EU (140.76%), LIFDC (56.37%) and NFIDC (0.88%); except – lower in Africa (42.99%), LDC (50.31%), LLDC (21.7%) and SIDS (35.47%). Phosphorus use in agriculture (%) is decreasing in the world (6.98%), Africa (0.25%), Americas (18%), North America (35.42%) and Europe (0.01%) than 2002 level. However, this value is still very high (65-99%) in most of these regions.

#### 3.1.6 Ecological footprint (EF)

Cropland EF per capita (gha) has been increased significantly in the word (14.87%), Africa (16.5%), Asia (55.22%), Europe (7.4%), north (8.89%) and South America (30.24%) and Oceania (3.21%), since 1961 level. Present level of cropland EF contributes a sizable portion to global average per capita ecological footprint PB, like – the word (33.35%), Africa (25.43%), Asia (30.31%), Europe (51.96%), North (65.63%) and South America (36.33%) and Oceania (33.59%) (Supplementary fig 7). The contribution of agriculture (i.e. cropland EF) to the respective total ecological footprint of consumption has increased in Africa (16.43%), Asia (22.26%), Europe (32.02%), north (40.79%) and South America (24.84%), Oceania (20.18%) and the world (21.39%) (Supplementary fig 8). Agriculture’s contribution (cropland EF) to Total EF used to be higher in countries Asia and Africa in 1961. Condition of countries of Asia has improved and of Africa has deteriorated. At the recent time too (2016), they hold the highest position than any other parts in the world.

**Fig 1:**
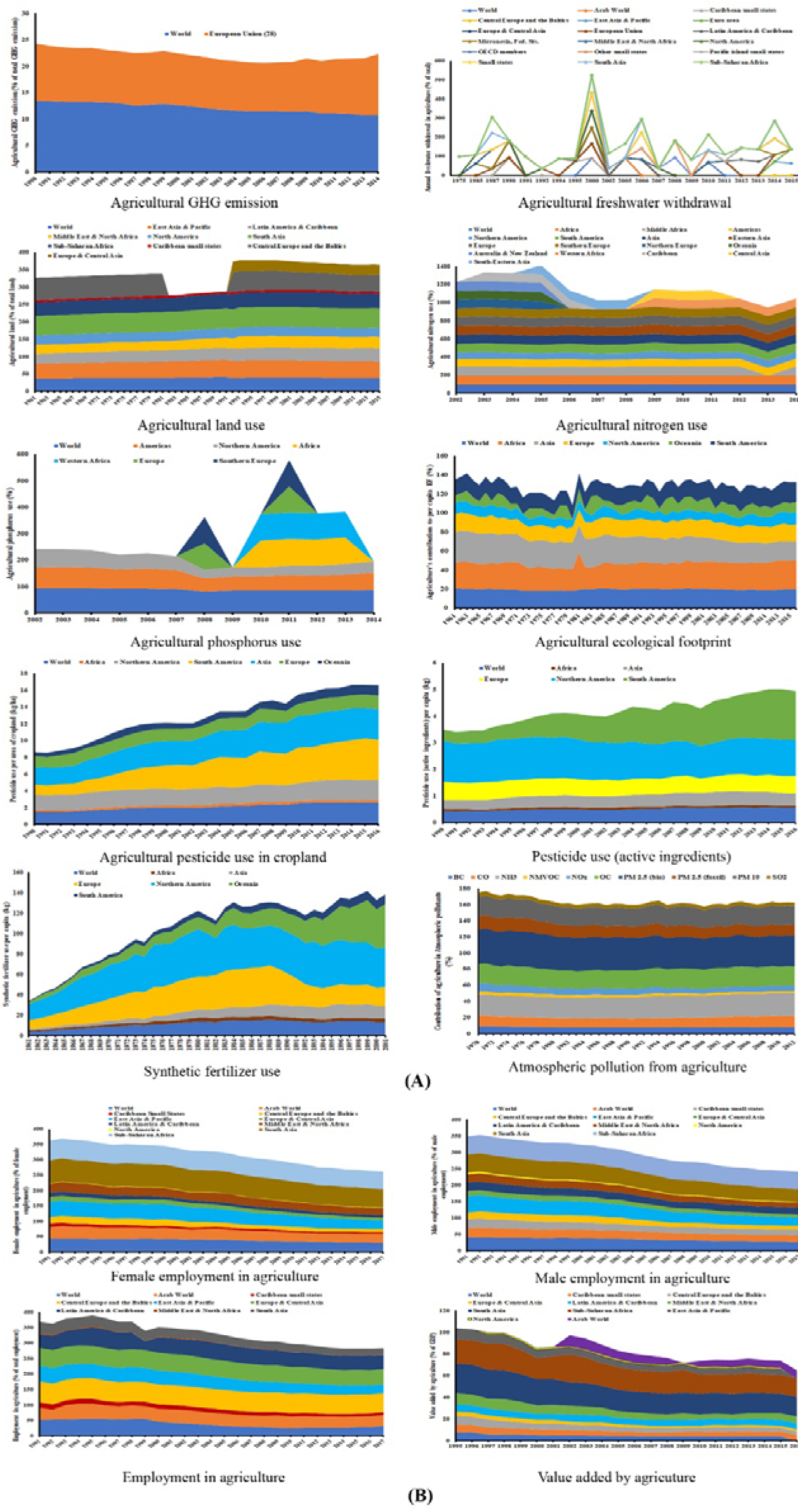
Trends of the contribution of global agriculture on (A) biophysical consumptions related to the planetary boundaries framework and (B) socioeconomic development.

**Fig 2:**
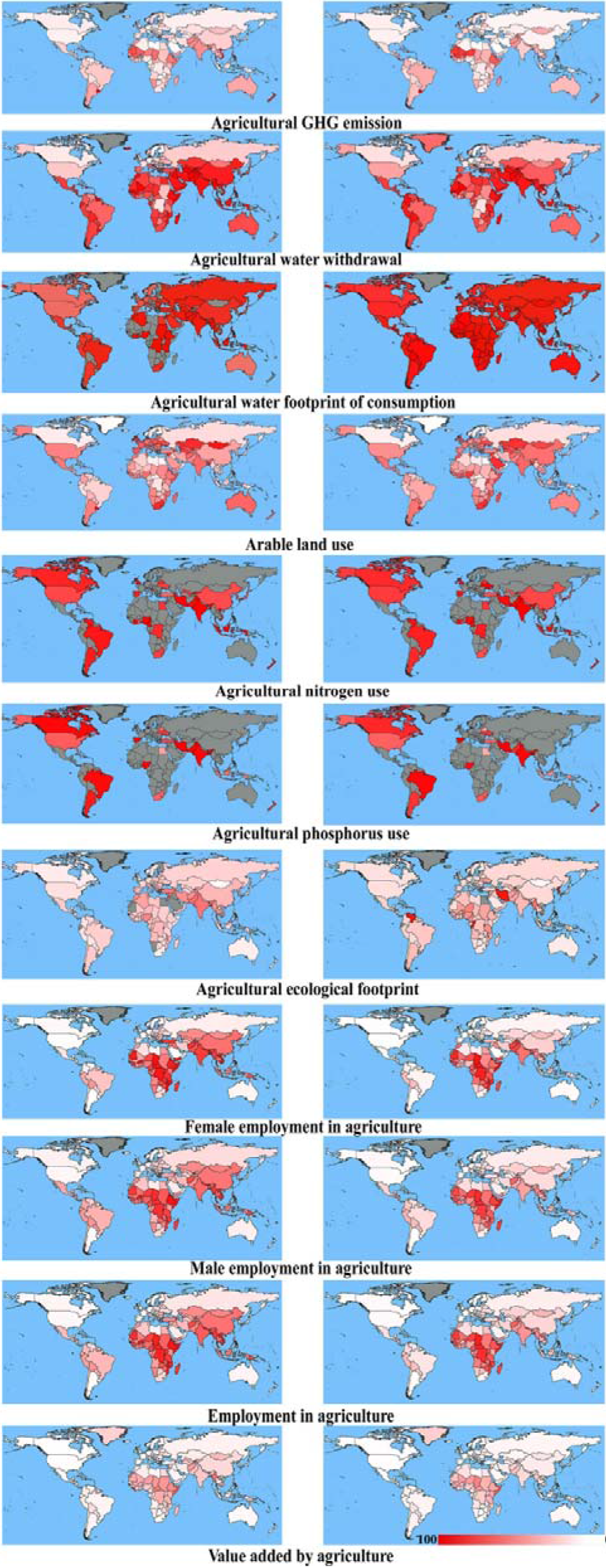
Changes in the contribution of agriculture in 7 dimensions of biophysical consumption related to planetary boundaries and 4 dimensions of socioeconomic development in countries of the world. A higher concentration of red indicates a higher contribution of agriculture (%) of a country. Seven dimensions of biophysical consumption are (1) Climate change, (2) Freshwater withdrawal, (3) Water footprint, (4) Land use, (5) Nitrogen use, (6) Phosphorus use, (7) Ecological footprint. Four dimensions of socioeconomic development are: (1) Female employment in agriculture, (2) Male employment in agriculture, (3) Employment in agriculture and (4) Value added by agriculture.

#### 3.1.7 Air pollution

Ammonia emission from agricultural sources was high in Europe, northern Asia and America in the 1970s. Now it’s high in the southeast and north Asia, few regions of South America, Europe and North America. Black carbon emission from agricultural sources was high south & southeast Asia, South Africa, South America and Oceania during the 1970s. Now its higher in South America and Oceania. Carbon monoxide emission from agricultural sources was high (the 1970s) in South Africa, south, southeast & West Asia and South America. But, now its higher in South America, Oceania, West & South Africa, Europe, south & southeast Asia. Agricultural emission of oxides of nitrogen was high in South America, Africa, south & southeast Asia. But presently its almost same except – decrease in south & southeast Asia. Non-methane volatile organic compounds emission from agricultural sources was very low except for a few countries in South America and Southeast Asia during the 1970s. But now it has increased in South America, Oceania, South Asia and parts Africa. Organic carbon emission from agricultural sources was high in central & South America, Oceania, middle east and southeast Asia during the 1970s. Presently, it’s very high in South America, Oceania, parts of south & southeast Asia; moderately high in north & West Asia, parts of Europe, North America, parts of West & South Africa. PM_2.5_ (bio) emission from agricultural sources was very high in North & South Africa, Europe, most of Asia; moderately high in South America, parts of Africa and Oceania during the 1970s. Now it’s even higher in both south & North America, most of Asia, Oceania and parts of Africa. PM_2.5_ (fossil) emission from agricultural sources was comparatively higher in parts of Africa, South America and South & Southeast Asia. Presently its decreased in all of those parts of the world. PM 10 from agricultural sources was moderately high in South America, almost all of Asia, Europe and Oceania. Now it has further increased, especially in South America, the Middle East and Oceania. SO_2_ emission from agricultural sources was very low all over the world, except – south & southeast Asia and parts of Africa during 1970s. But, now its slightly high in parts of South America, Africa, Europe, Southeast Asia and Oceania. At the global average level, the contribution of agriculture has decreased in all the of these air pollutants (except - increase in CO, NH3 and OC).

**Fig 3:**
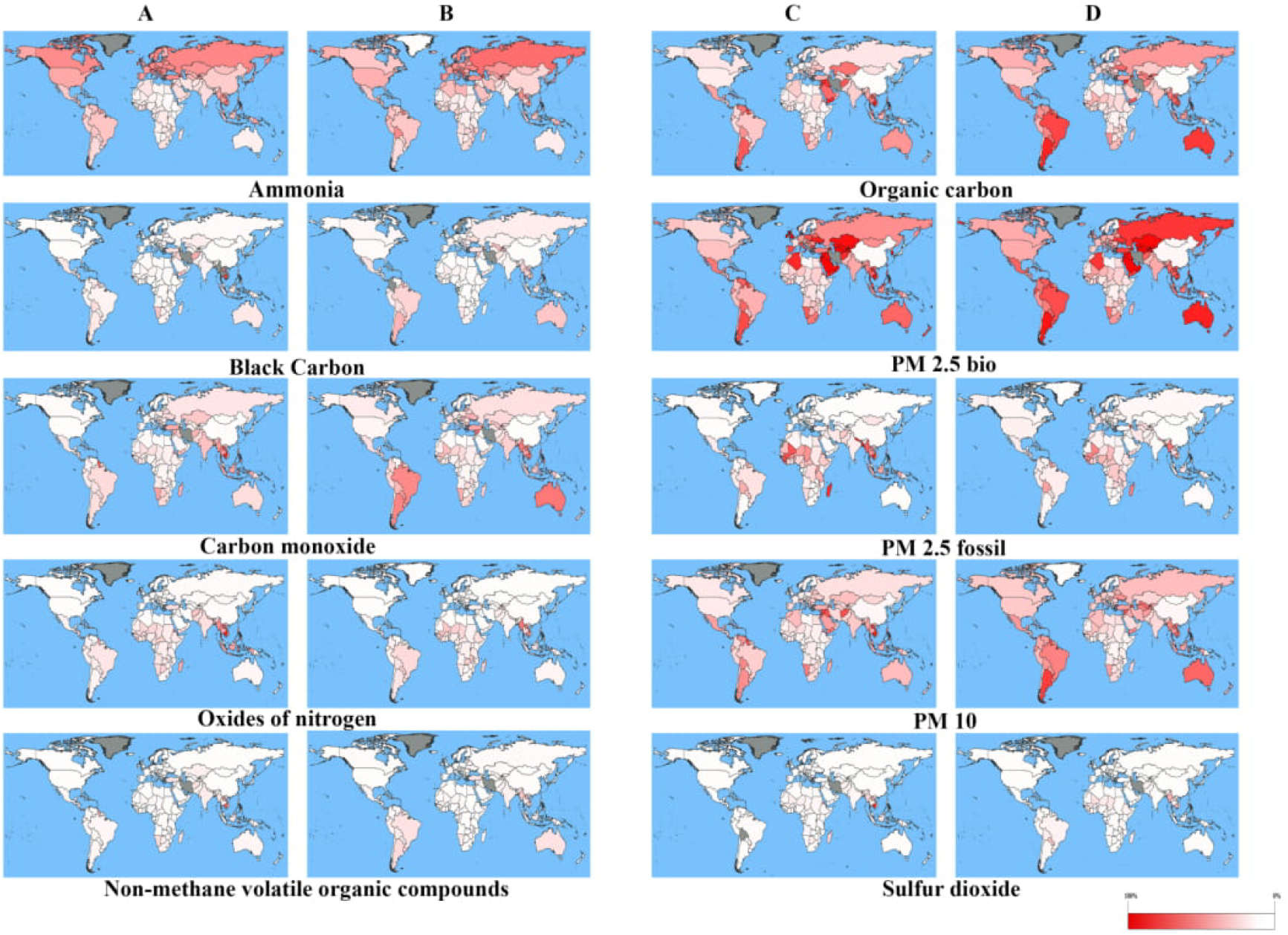
Changes in the contribution of agriculture in 10 indicators of atmospheric pollution related to planetary boundaries in countries of the world. A higher concentration of red indicates a higher contribution of agriculture (%) of a country. (A and C) represents the past level of indicators and (B and D) represents the most recent available values of respective indicators.

#### 3.1.8 Novel entities

Synthetic fertilizer use (kg/capita) has increased at a very high rate in the world (251.46%), Africa (145.93%), Asia (920.03%), Europe (106.98%), north (159.46%) and South America (702.21%), Oceania (1564.69%) since 1961 level. This has also increased in a similar way in other national groups, as – EU (107.63%), LDC (640.81%), LLDC (648.57%), SIDS (50.94%), LIFDC (1225.81%) and NFIDC (333.46%) (Supplementary fig 9).

Use of active ingredients of pesticides (kg/capita) have increased in the world (27.65%), Asia (38.8%), Oceania (76.99%) and South America (334.62%); but decreased in Africa (17.57%), Europe (3.6%) and North America (12.28%), since 1990 level (Supplementary fig 10). Agricultural pesticide uses per area of cropland (kg/ha) have increase relatively more in the world (41.63%), south America (76.49%), Asia (41.44%) and Oceania (64.1%), than Africa (3.22%), north America (24.37%) and Europe (19.76%) from 1990 level. In Africa, it has increased more in middle, south and western part (20, 39.25 & 50%, respectively), but decreased in east & northern part (9.52 & 2.74%). In America, both central part and Caribbean has increased (45.45 & 17.04%). In Asia, eastern and western part has increased (51.69 & 43.12%), but southern, south-eastern and central part has decreased (30, 2.25 & 21.82%, respectively). In Europe, eastern and southern part have increased (38.27 & 14.97%), whereas northern and western part have decreased (43.65 & 9.72%). In Oceania, Australia and New Zealand & Melanesia have increased (64.7 & 44.93%), but Polynesia have decreased (67.35%). Among other groups, LDC, LLDC, SIDS and NFIDC have increased (78.26, 58.93,17.42 & 37.78%, respectively), whereas EU & LIFDC have decreased (3.18 & 19.05%) (Supplementary fig 11).

**Fig 4:**
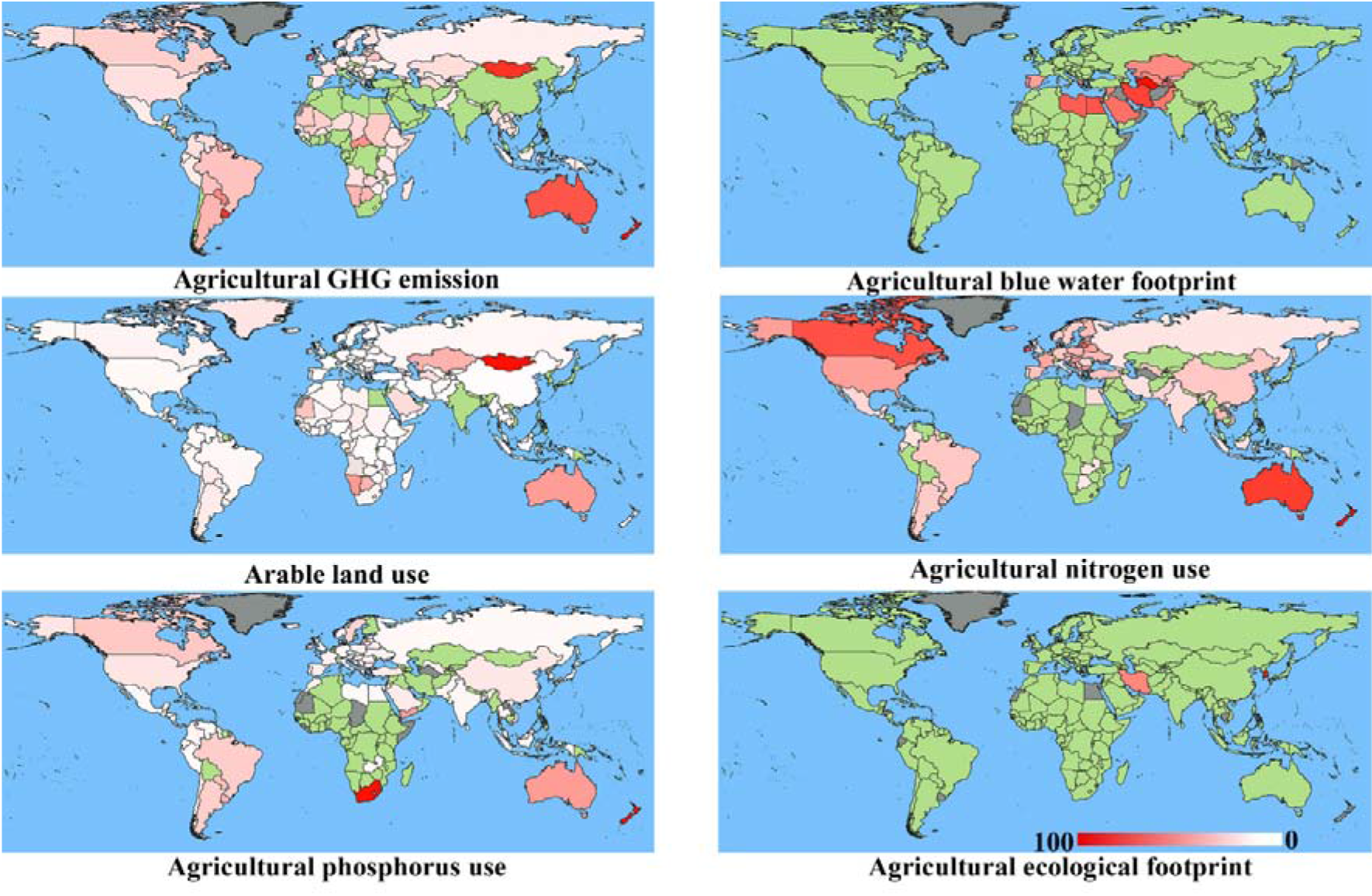
Recent scenario of countries in the world on six dimensions of biophysical consumption related to safe operating space of agriculture (i.e. planetary boundaries framework). These dimensions are – climate change, blue water footprint, land use, nitrogen use, phosphorus use and ecological footprint. Green and red colours indicate countries which are within or exceeded safe limits of respective dimensions of planetary boundaries. A higher concentration of red indicates how far away the country is from a safe limit.

### 3.2 Future scenario

According to World Urbanization Prospects (WUP 2018) projection of population at global scale, use of various dimensions of planetary boundaries are going to increase in 2050 as follows – climate change (25.2-37.7%), arable land use (12.72-65.08%), ecological footprint (13.29-33.24%), nitrogen use (14.62-25.82%), phosphorus use (10.47-24.82%), synthetic fertilizer use (24.44%), pesticide use (23.5-26.33%). At highest per capita level, for all the SSPs it will increase 28.04-38.81% (climate change), 59.67-65.7% (arable land use), 16.76-37.23% (ecological footprint), 14.31-27.14% (nitrogen use), 13.16-26.16% (phosphorus use), 37.38-46.76% (synthetic fertilizer use) and 14.9-27.64% (pesticide use). However, at the lowest per capita level, it is possible to check this increase, at least up to a certain degree. For example, for all the SSPs at lowest level these will increase as – 13.65-26.58% (climate change), 12.72-25.79% (arable land use), 1.37% decrease to 15.99% increase (ecological footprint), 1.38-16.14% (nitrogen use), 3.41% decrease to 12.07% increase (phosphorus use), 119.85-158.56% decrease (synthetic fertilizer use) and 15.95% decrease to 1.41% increase (pesticide use).

**Fig 5:**
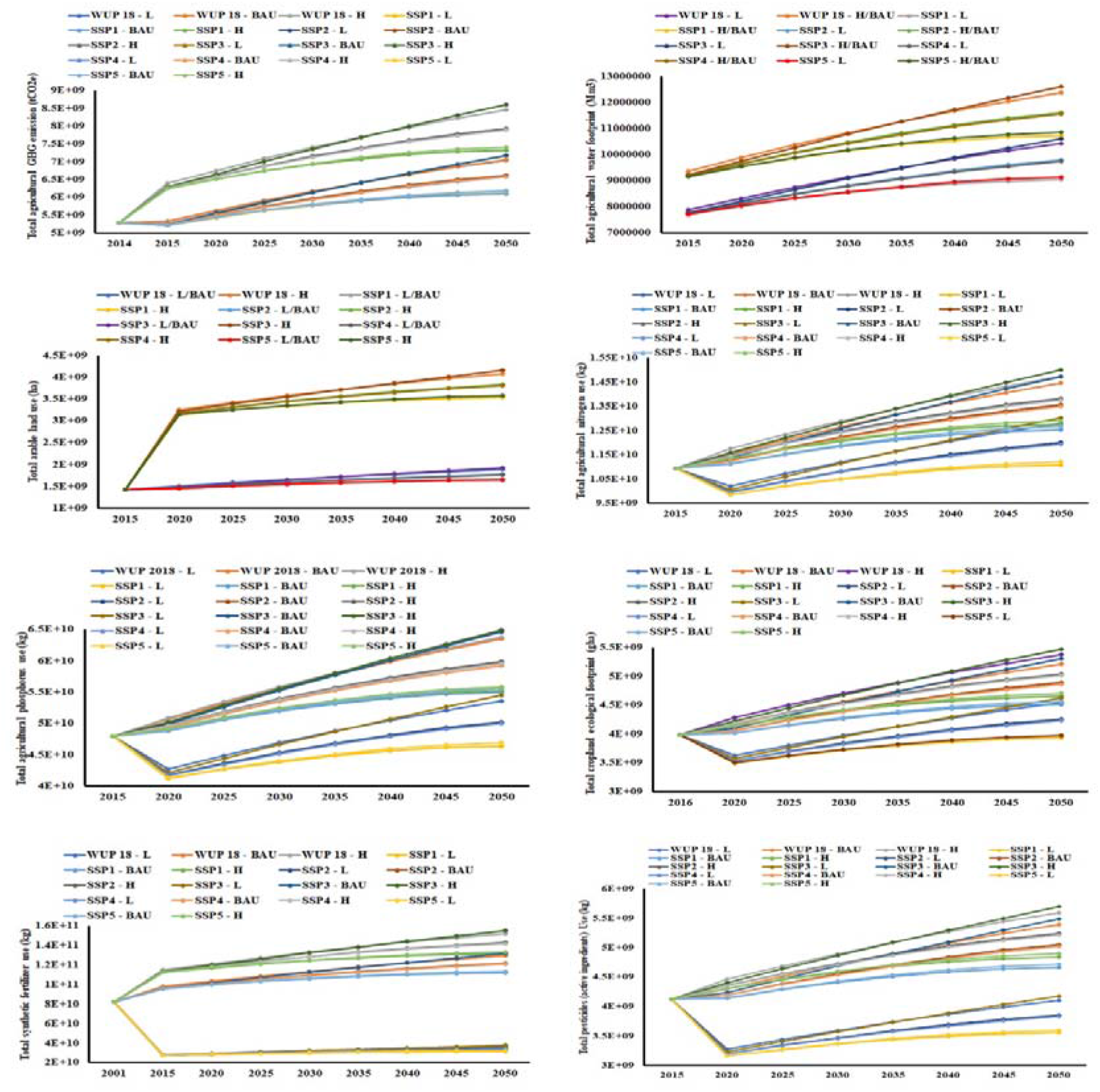
Projection of 8 dimensions of biophysical resource consumption in global agriculture in future up to 2050. The 6 scenarios are – 5 SSPs (shared socioeconomic pathways) and 1 UN (World Urbanization Projection, 2018). It is projected on 3 levels of consumption: lowest, BAU (based on data availability) and highest.

If the most efficient path of population growth is not chosen over the opposite, biophysical safe limits, related to agriculture, can decrease by almost 55% (climate change), 300% (freshwater use), 50-55% (arable land use), 180% (nitrogen use), 265% (phosphorus use) and 20% (ecological footprint) in 2050.

**Fig 6:**
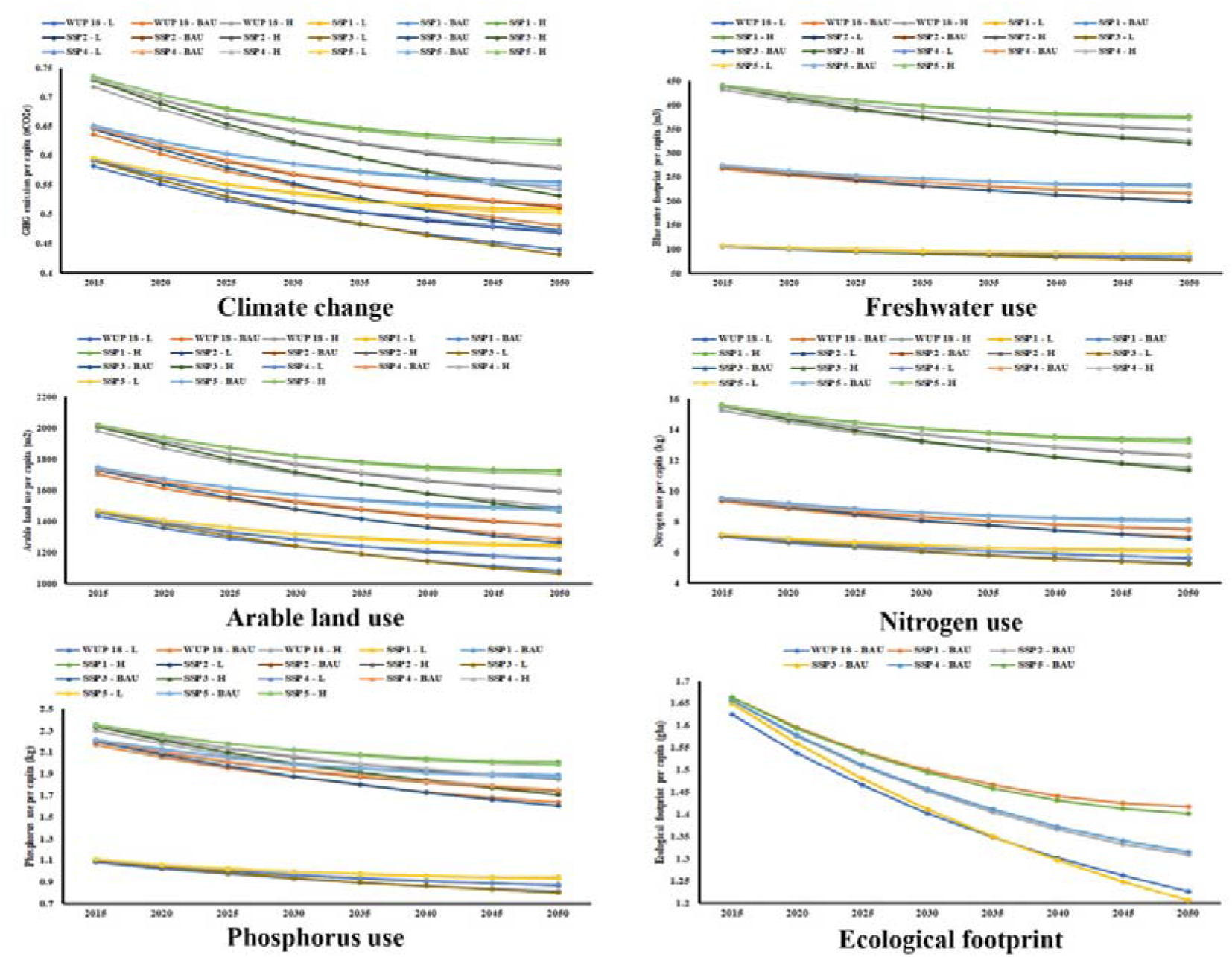
Projection of availability of biophysical resources for agriculture up to 2050. Based on 5 SSPs and 1 WUP 2018 projection. The 6 scenarios are – 5 SSPs (shared socioeconomic pathways) and 1 UN (World Urbanization Projection, 2018). It is projected on 3 levels of consumption: lowest, BAU (based on latest data availability) and highest.

### 3.3 Doughnut economy of agriculture

At the global level, with the increase of GDP per capita (current US$), agricultural GHG emission, arable land use and freshwater use have decreased gradually. However, the remaining 5 dimensions (i.e. ecological footprint, nitrogen and phosphorus use, synthetic fertilizer and pesticide use) have increased (Supplementary fig 12). Also, among the atmospheric pollutants, ammonia and PM_2.5_ (fossil) emissions have increased (Supplementary fig 13). With the global level increase of GNI per capita (constant 2010 US$), similar results are found (Supplementary fig 14, 15).

Female employment in agriculture (% of total female employment) has decreased overall in 27 years (1991-2017). It has decreased less in Caribbean Small States, Latin America & Caribbean, Middle East & North Africa, North America and Sub-Saharan Africa (6.78, 4.58, 5.86, 0.67 and 6.24%, respectively); it has decreased more in the World, Arab World, Central Europe and the Baltics, East Asia & Pacific, Europe & Central Asia and South Asia (12.57, 9.16, 10.64, 21.5, 7.18 and 14.09%, respectively). Among the other regions, it has also decreased to a varying degree. This has decreased less in the Euro area, HIPC, OECD Members and PISS (4.75, 2.79, 4.82 and 0.88%, respectively); and has decreased more in EU, FACCs, LDCs, Other Small States and Small States (5.8, 9.13, 8.12, 9.92 and 8.06%, respectively). In all of the income groups, it has decreased, as - HI, L&MI, LMI, MI, UMI (3.59, 15.79%, 18.15, 18.83 and 20.7%, respectively), except – LI group (0.008% increase).

In 27 years (1991-2017), male employment in agriculture (% of total female employment) has also decreased. It has decreased less in Europe & Central Asia, Latin America & Caribbean, North America and Sub-Saharan Africa (5.61, 5.31, 2.16 and 2.35%, respectively); it has decreased more in the World, Arab World, Caribbean small states, Central Europe and the Baltic, East Asia & Pacific, Middle East & North Africa and South Asia (13.46, 8.74, 9.71, 9.08, 25, 9.07 and 17.51%, respectively). Among the other regions, it has also decreased; less in Euro area, EU and OECD members (3.67, 4.5 and 4.41%, respectively), more in FACCs, LDCs, Other small states, PISS and Small states (7.63, 6.87, 13.69, 7.43 and 12.67%, respectively), except in HIPCs (0.1% increase). In all of the income groups, it has decreased, as - HI, L&MI, LMI, MI, UMI (3.54, 17.13, 16.9, 19.36 and 23.08%, respectively), except – LI (2.02% increase).

In 27 years (1991-2017), employment in agricultural sector has decreased more in the World, East Asia & Baltics, and South Asia (19.1, 23.46 and 15.7% respectively); lesser level of decrease is found in Arab world, Caribbean, central Europe & Baltics, Europe & Central Asia, Middle East & North Africa (4.09, 9, 5.29, 8.18 and 1.6%, respectively). However, it has only increased slightly (2.7%) in Latin America & the Caribbean. Likewise, employment in the agriculture sector has also decreased in the Euro area, EU, FACCs, HIPCs, LDCs, OECD members, Other small states and Small states (5.12, 3.1, 1.1, 8.5, 7.48, 12.32, 12.8 and 3.9%, respectively); increased in PISS (23.3%). This agriculture’s share in employment has also decreased all the categories of income groups, as - HI, L&MI, LI, LMI, MI (3.59, 22.98, 7, 2.1 and 0.1%, respectively); increased only in UMI (1.1%).

Value added by agriculture (% of GDP) has decreased in 22 years (1995-2016) in all of the regions in the world. It has decreased less in Caribbean small states, Europe & Central Asia, Latin America & Caribbean and North America (1.88, 1.95, 1 & 0.33%, respectively); and decreased more in World, Central Europe and the Baltics, Middle East & North Africa, South Asia, Sub-Saharan Africa, East Asia & Pacific and Arab World (4.07, 4.74, 4.08, 9.27, 5.21, 4.63, & 3.04%, respectively). In 16 years (2000-2016), this has decreased less in the Euro area, EU, OECD members, Other small states & Small states (0.78, 0.82, 0.54, 1.53 & 1.52%, respectively); and decreased more in HIPCs, LDCs, PISS and FACCs (5.07, 4.85, 7.21 and 6.06%, respectively). In 31 years (1986-2016), value-added by agriculture (% of GDP) has decreased in all of the income groups, as – L&MI, LI, LMI, MI, UMI & HI (11.92, 11.08, 12.29, 11.96, 12.13 & 0.79%, respectively) (Supplementary fig 16).

The relationship among the 22 indicators, through correlation table (see Supplementary table) and correlogram (Supplementary fig 17) has also been shown.

## 4. Discussion

At present day, the industry doesn’t have the capacity to replace agriculture (via synthesized food etc.) and sufficiently feed billions of populations. Also, giving priority on biodiversity alone over agriculture would again fail to sustain the populace. Therefore, agriculture is irreplaceable at this present state of advancement.

There are few things, related to biophysical consumption of agriculture, to be inferred from this work. First, countries, where resource consumption (GHG emission, water use, water footprint, land use, nitrogen and phosphorus use, cropland EF) from agriculture has increased over the years, should change the process, following the example of those countries where the opposite has happened. Agriculture should not become a ‘necessary evil’, especially when it has to feed billions of people both now and in future. Second, there should be a balance in what is being invested into agriculture (biophysical resources, manual labour, time etc.) and what is the outcome from it (food, nutrition, social and economic development etc). Third, we should always try to gain more out of agriculture without or keeping input to agriculture and pollutions generated from it (greywater, soil erosion, agrobiodiversity loss, leaching of N and P manures) in check. In this way, we can restructure agriculture towards a sustainable and smart process in accordance with the necessities of present times and upcoming future. Fourth, the contribution of agriculture in the emission of atmospheric pollutants should be decreased, especially – ammonia, carbon monoxide, organic carbon, PM_2.5_ (bio) and PM_10_. Fifth, crops need to be redistributed according to distribution and abundance of the ingredients of agricultural products, soil features, hydrogeology etc (Davis et al., 2017, Rosa et al., 2018). Continued stress on monoculture (due to cultural, dietary preference etc.) would result in a negative feedback effect sooner or later, with adverse consequences on both environment and economy. Sixth, the status of economic debit or credit should not be the driving force of trade of agricultural and agriculture-derived commodities for any region or country. This can distort biophysical resources equity.

There are a few points, related to socioeconomy of agriculture, to be inferred from this work. First, from the perspective of gender equality, very few countries have achieved gender-wise equal employment ratio at the almost same level. This should be considered a desired goal for all the remaining countries with existing gender disparity related to the agriculture sector. Second, employment generation is not good enough from agriculture in some developing countries whereas biophysical resource depletion is fairly enough. These require an increase and decrease, respectively. These countries might learn from those where employment generation in the agriculture sector is enough with minimal resource consumption. Third, from an economic viewpoint, very few countries presently have a significant contribution to agriculture in their GDP whilst reserving resource consumption at a low level. This should be practised in all the other countries too so that agriculture can become an economically productive and efficient sector. Fourth, Social and economic development generated from agriculture should be given equal significance, especially by governments and research community, with biophysical resource consumption in agriculture, to make agriculture sustainable, in totality.

There are few things, related to the safe operating space of agriculture, to be inferred from this work. First, among the 6 biophysical consumption indicators used in this work, 4 (viz. climate change, land use, nitrogen and phosphorus use) needs immediate attention in most of the parts of the world. Second, if the ecological footprint is to be used as one of the biophysical consumption indicators related to planetary safe space concept, sector-wise ecological footprint allocations are necessary for improved assessment of resource consumption on per capita basis. Third, more detailed work on sector-specific safe limits of different types of water (blue, green or grey) is needed. Fourth, as per our understanding from this work, overpopulation could be one of the reasons for being under per capita safe limits of land, nitrogen and phosphorus use PB in countries of Africa and Asia. Their performance should not be interpreted as a good condition.

There are a few limitations of this work which also provides scope for work in this dimension in future. First, among the three major areas of consumption, only the safe operating space related to agriculture has been formulated so far. Formulation of sector-specific safe limits of biophysical consumption for the other two major sectors (industrial and household consumption) is necessary to understand their inter-sector comparison and interactions. Second, if more comprehensive data on other aspects related to socioeconomy of data was available, it would have been possible to completely analyse agriculture from the perspective of doughnut economy. This would result in a more holistic understanding of global agriculture. Third, hierarchical contribution of different agricultural products or agricultural systems is desired to make our understanding more specific. Fourth, it is important that the safe operating space of ‘introduction of novel entities’ should be established, which has special importance in the agricultural sector. Fifth, long term local to global scale multi-indicator data about the effect of agriculture on biodiversity loss at the group level (both animal and plant) should be available. Sixth, water has one of the most significant impacts on agriculture. However, sector-specific safe limits of various types of water (blue, green and grey water) are not available yet. Seventh, though ecological footprint has been used as a good indicator of biophysical consumption in the planetary boundaries framework, sector-specific allocation of safe limits (in gha) is necessary to use it in more specific purposes. Eighth, spatio-temporal data of long-time span at the local level is needed to integrate this planetary boundary framework of agriculture into policymaking from local to national level of governance.

This work results in a global exhaustive overview of safe and just operating space and agriculture up to country-level. We have also tried to find the conflicts and deficiencies where further works need to be done to integrate it in policy framing in future.

## Acknowledgement

This study was supported by FRPDF scheme of Presidency University, Kolkata. We would like to thank Agniswar Chakraborty, Department of Computer Science, Jadavpur University (India) for his kind assistance during the preparation of correlogram.

## Footnote

Works done in this paper is part of a doctoral thesis to be submitted in partial fulfilment of the requirements of a degree of Doctor of Philosophy from Presidency University, Kolkata.

