## Supplementary figure for "Analysing the safe and just operating space of agriculture in the world: past, present and future"

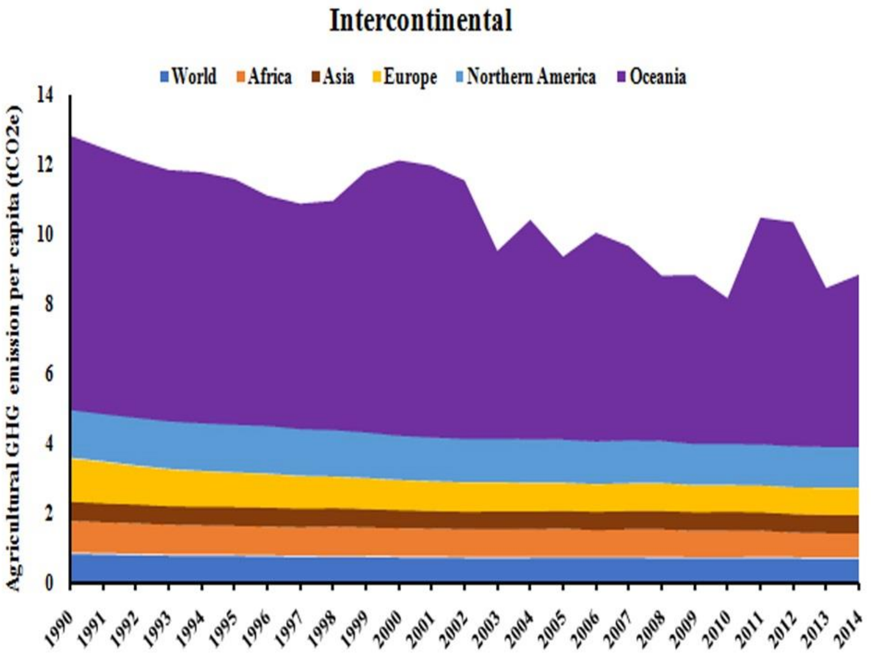


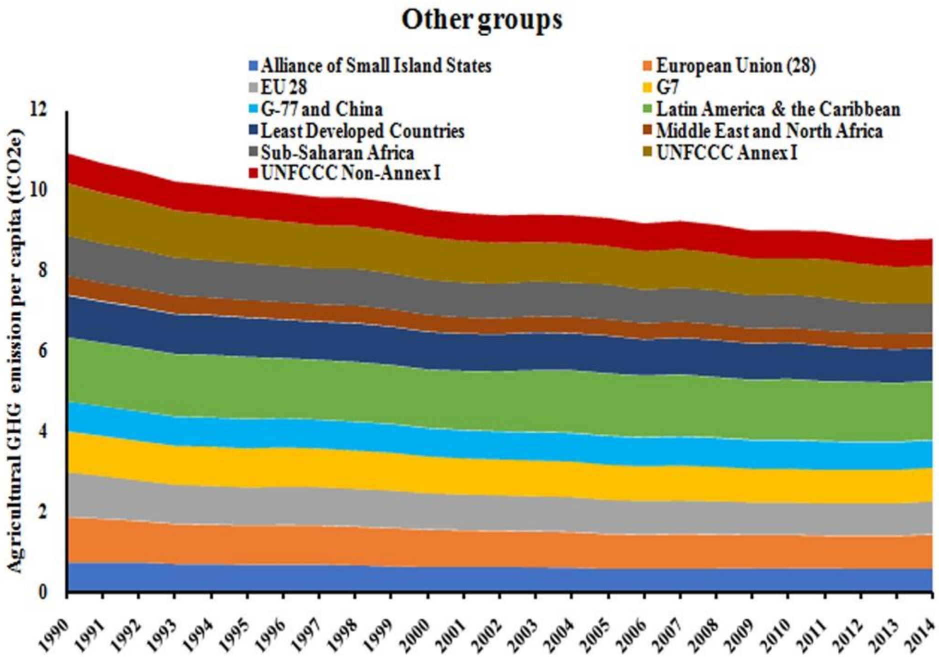


**Fig 1: Trends of per capita agricultural GHG emission in regions of the World.**

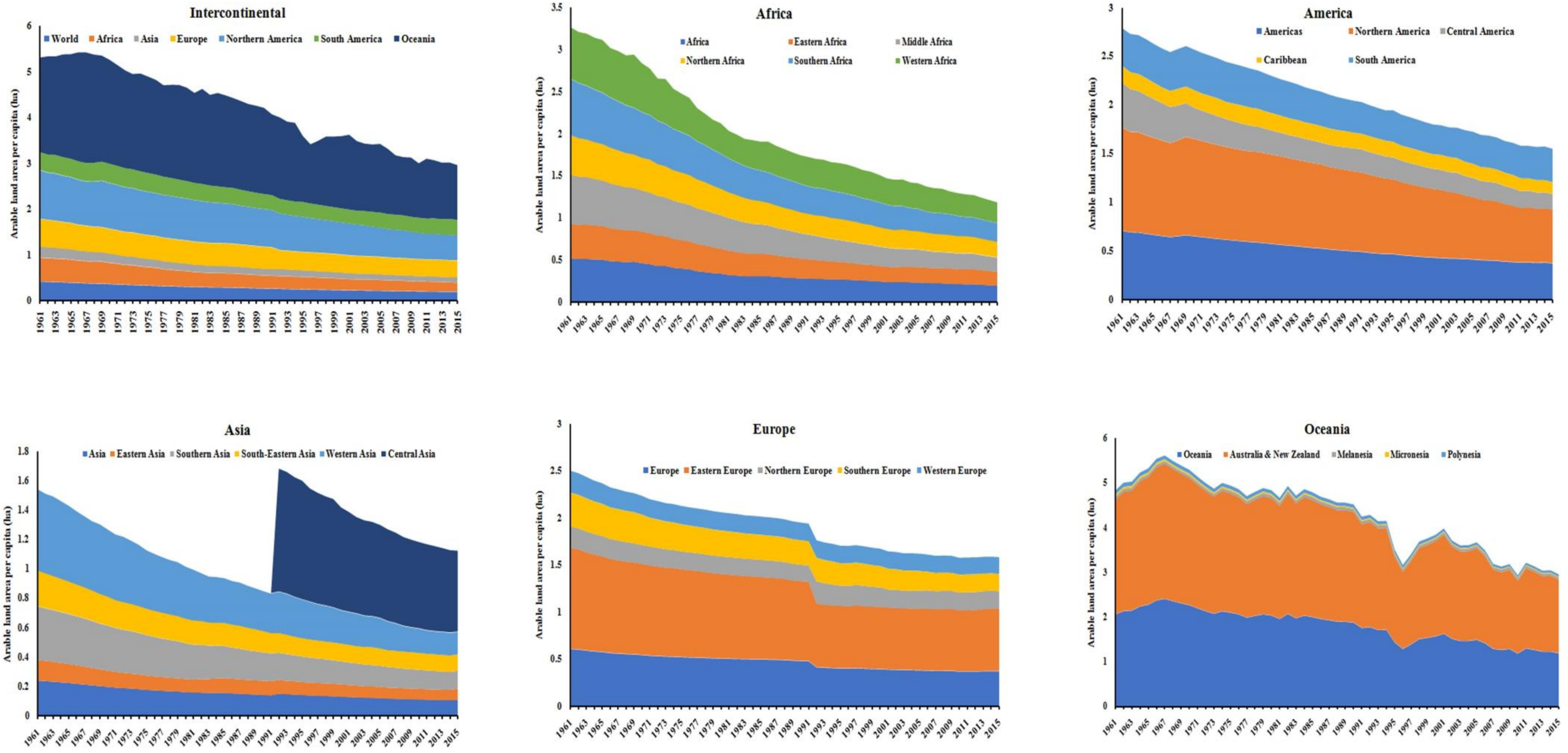


**Fig 2: Trends of per capita agricultural land use in regions of the World.**

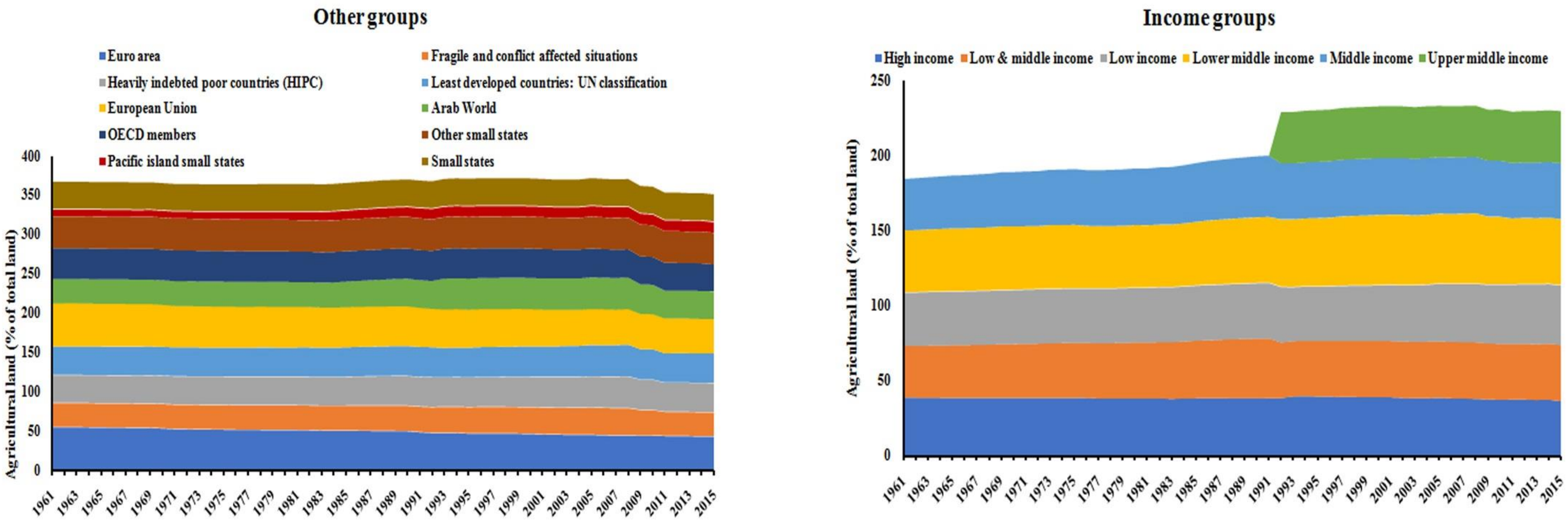


**Fig 3: Trends of contribution of agriculture (%) in land use in regions of the World.**

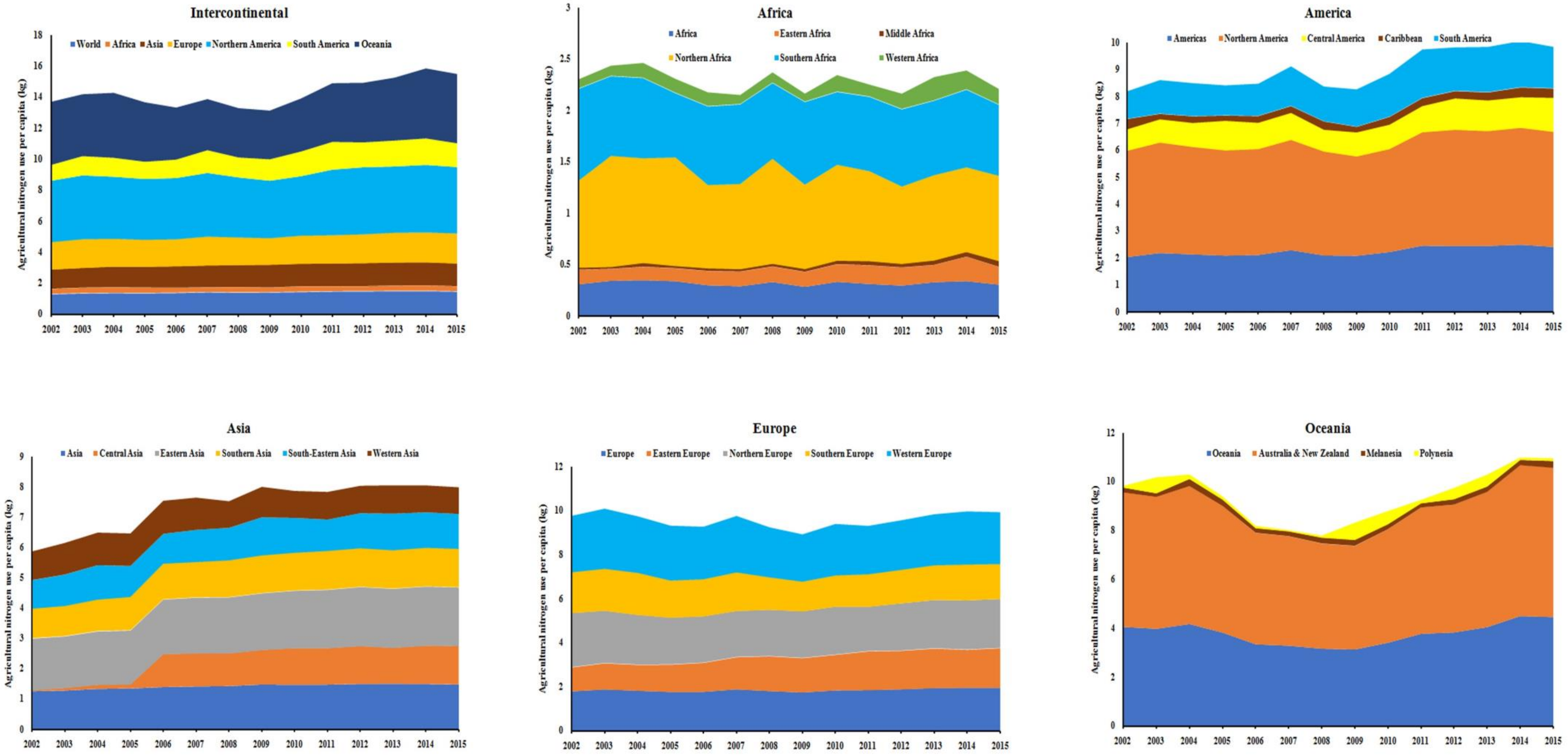


**Fig 4: Trends of per capita agricultural nitrogen use in regions of the World.**

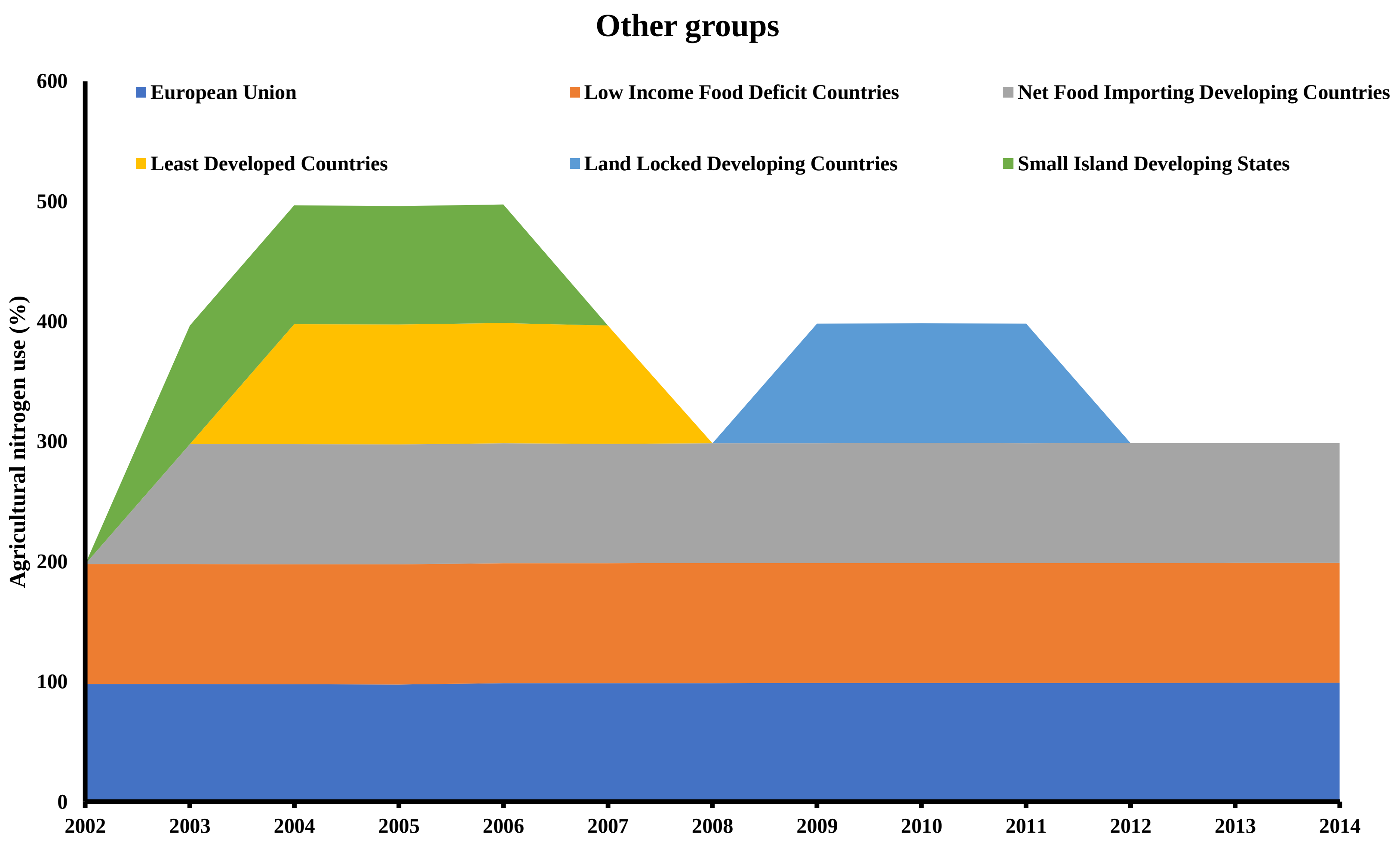


**Fig 5: Trends of contribution of agriculture (%) in nitrogen use in regions of the World.**

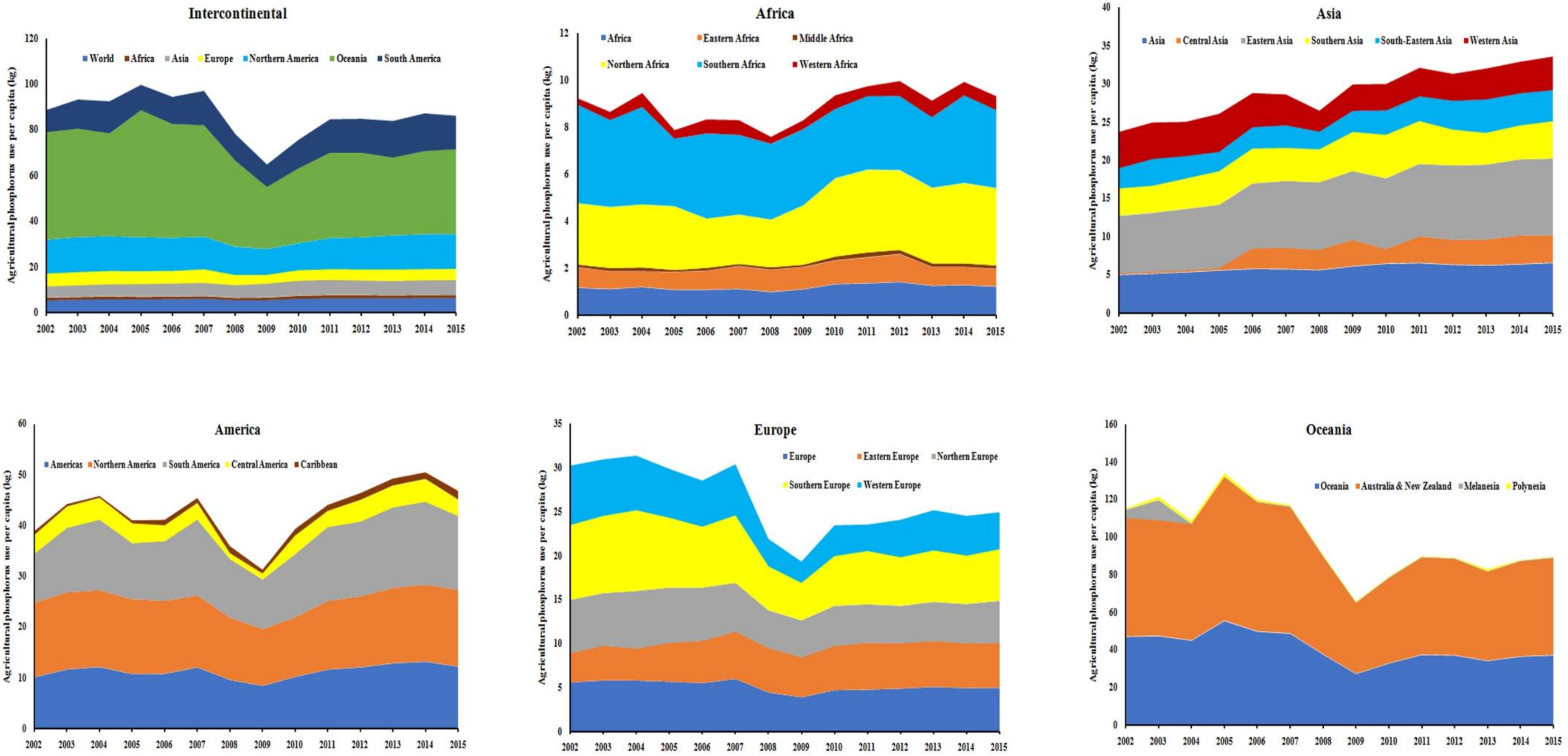


**Fig 6: Trends of per capita agricultural phosphorus use in regions of the World.**

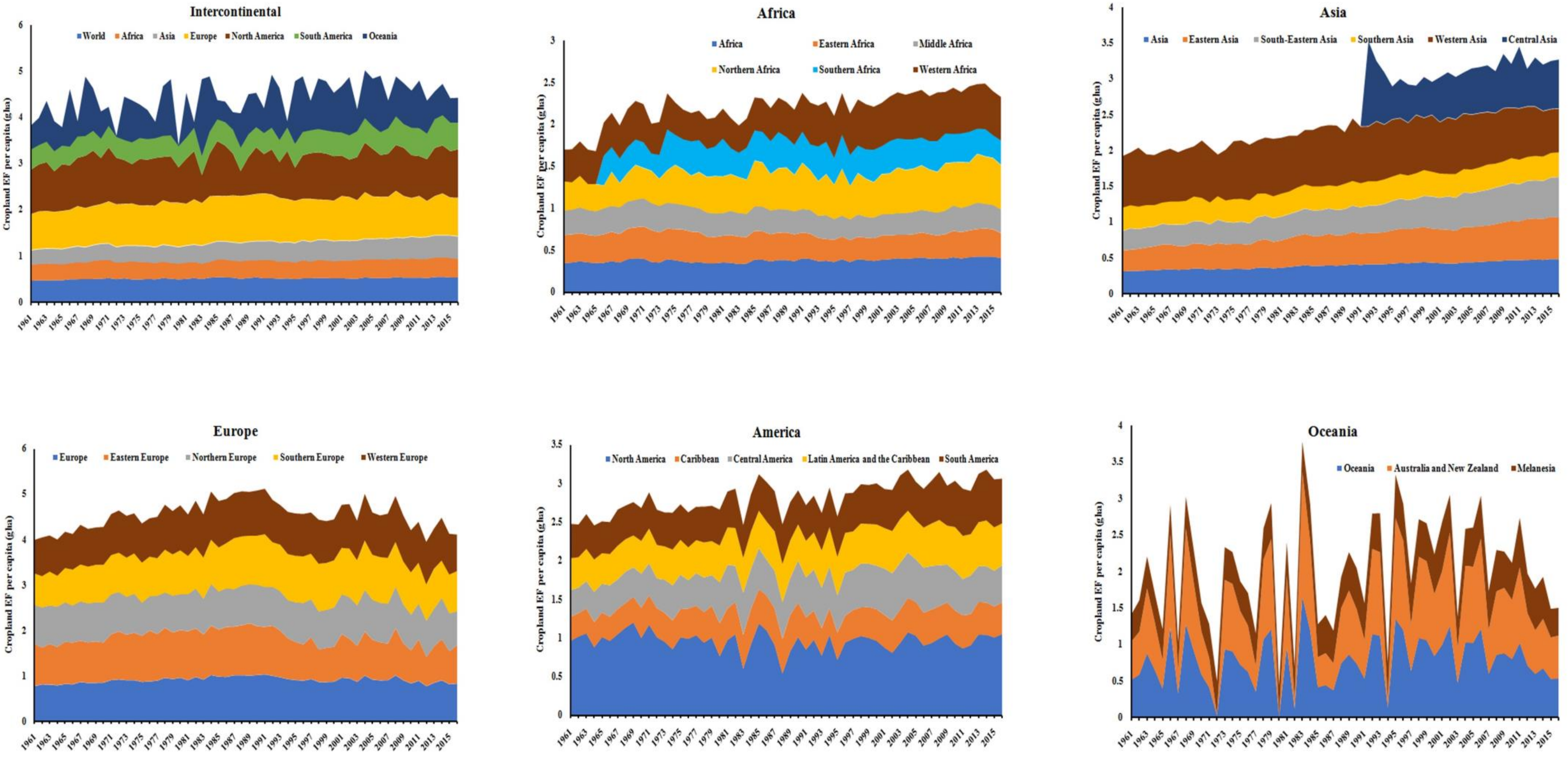


**Fig 7: Trends of per capita agricultural ecological footprint in regions of the World.**

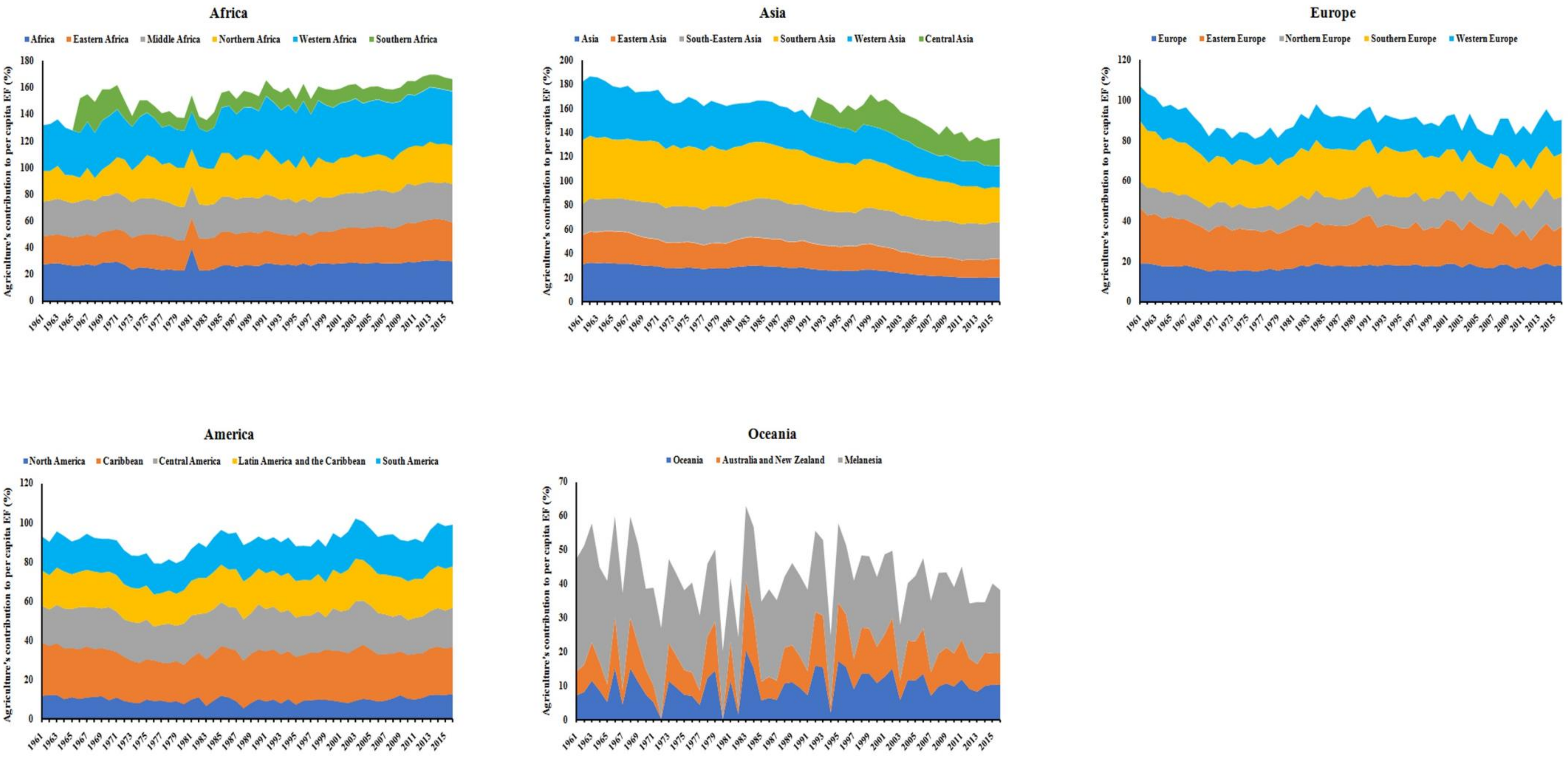


**Fig 8: Trends of contribution of agriculture (%) in ecological footprint in regions of the World.**

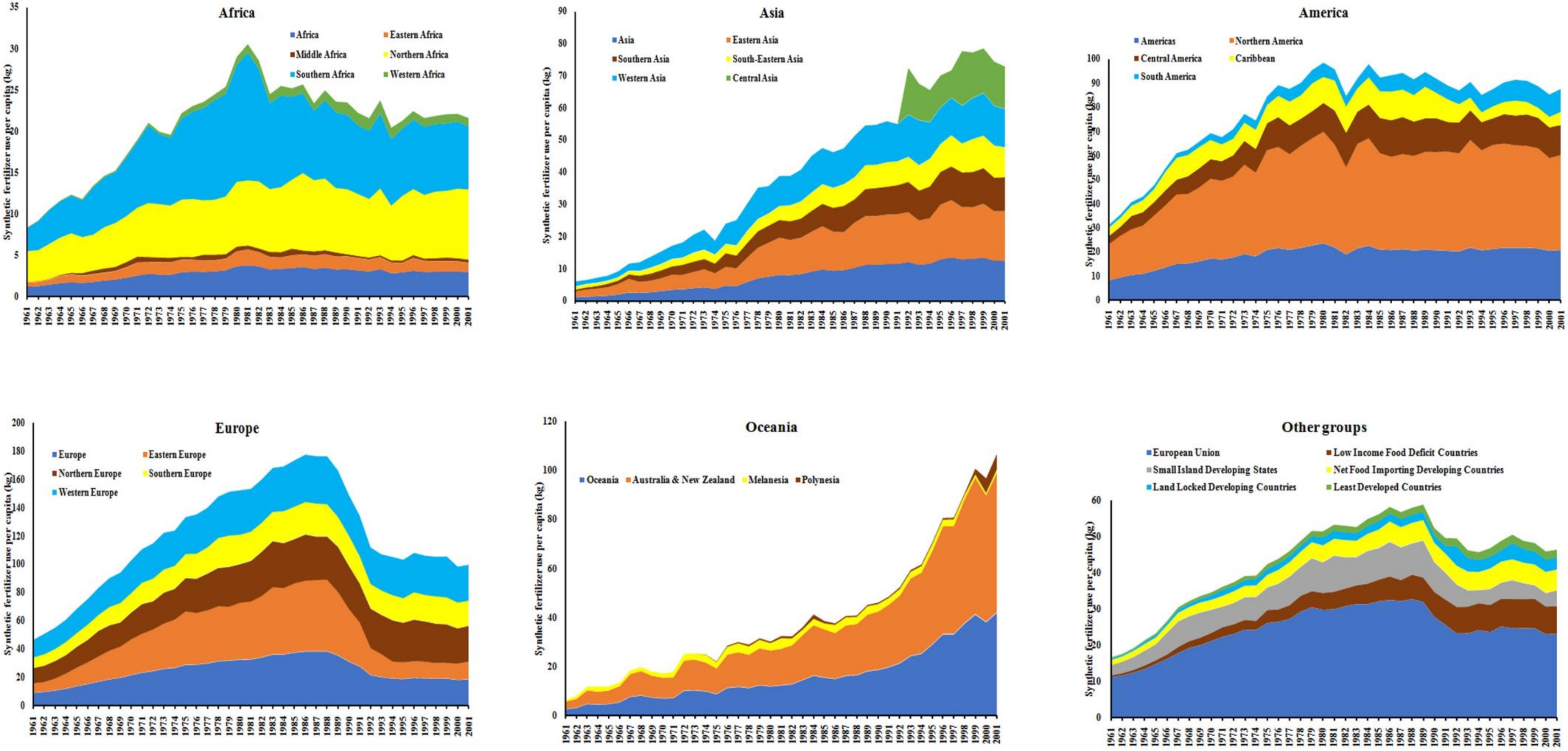


**Fig 9: Trends of per capita agricultural synthetic fertilizer use in regions of the World.**

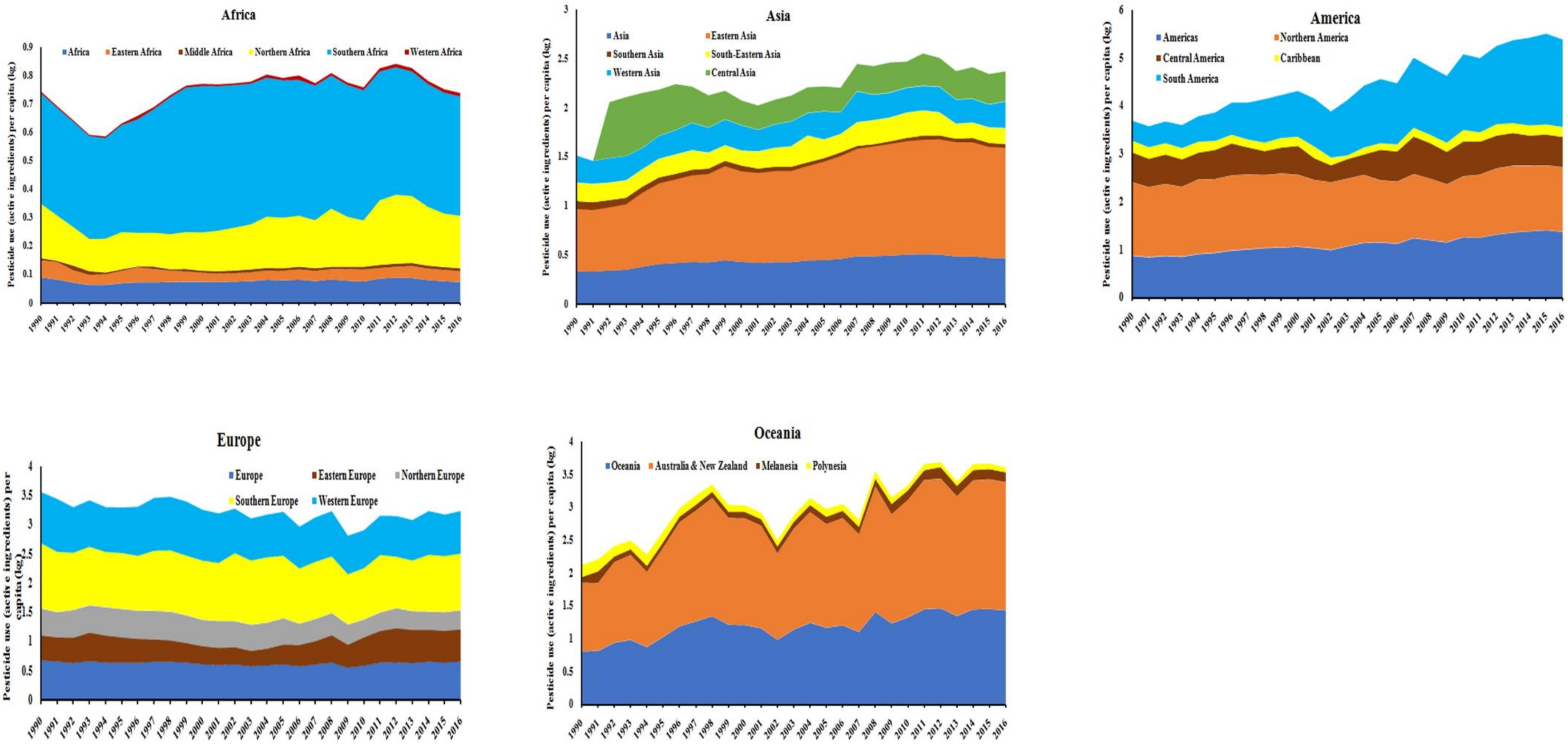


**Fig 10: Trends of per capita agricultural pesticide use (active ingredients) in regions of the World.**

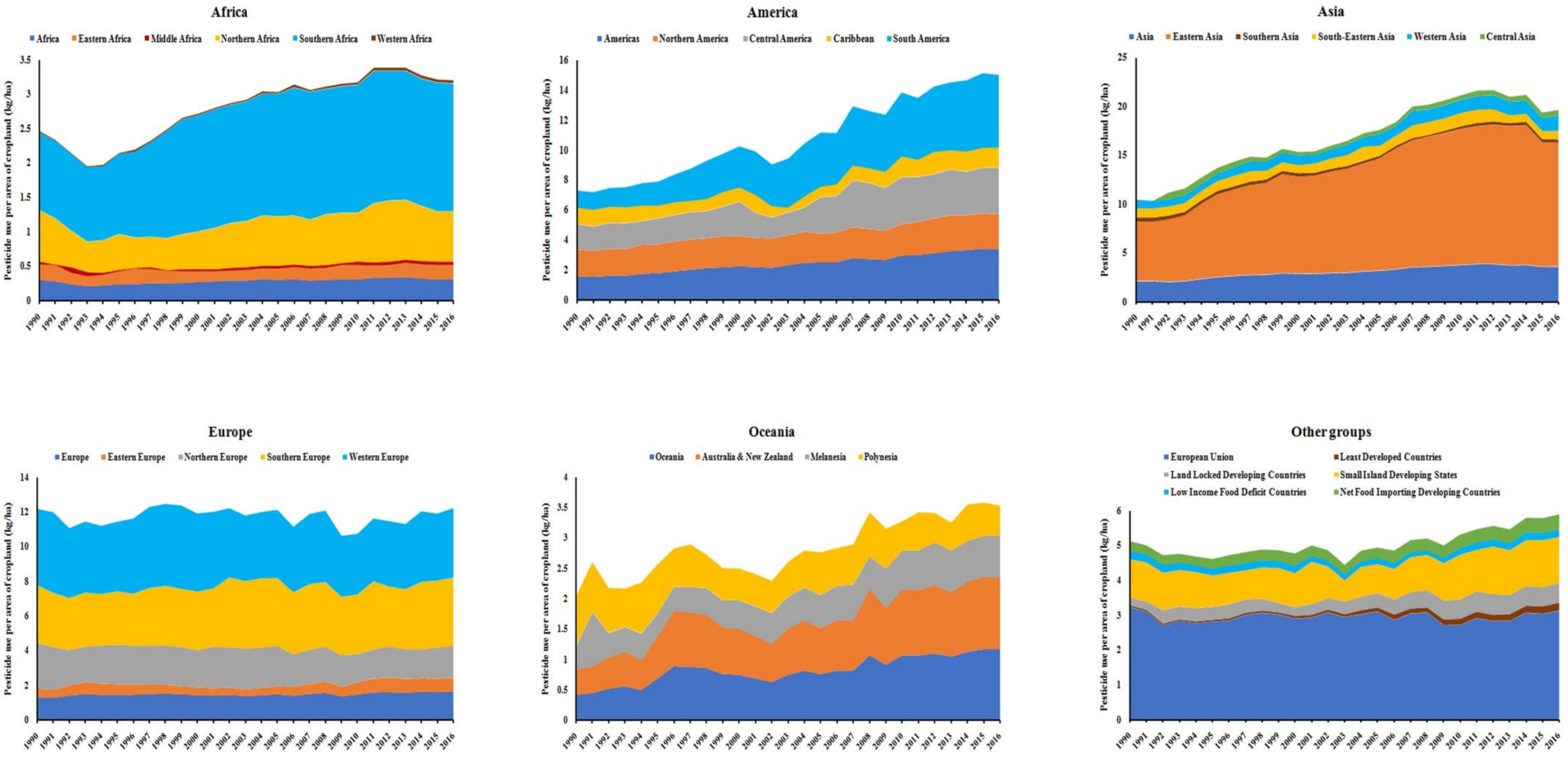


**Fig 11: Trends of per capita agricultural pesticide use in cropland in regions of the World.**

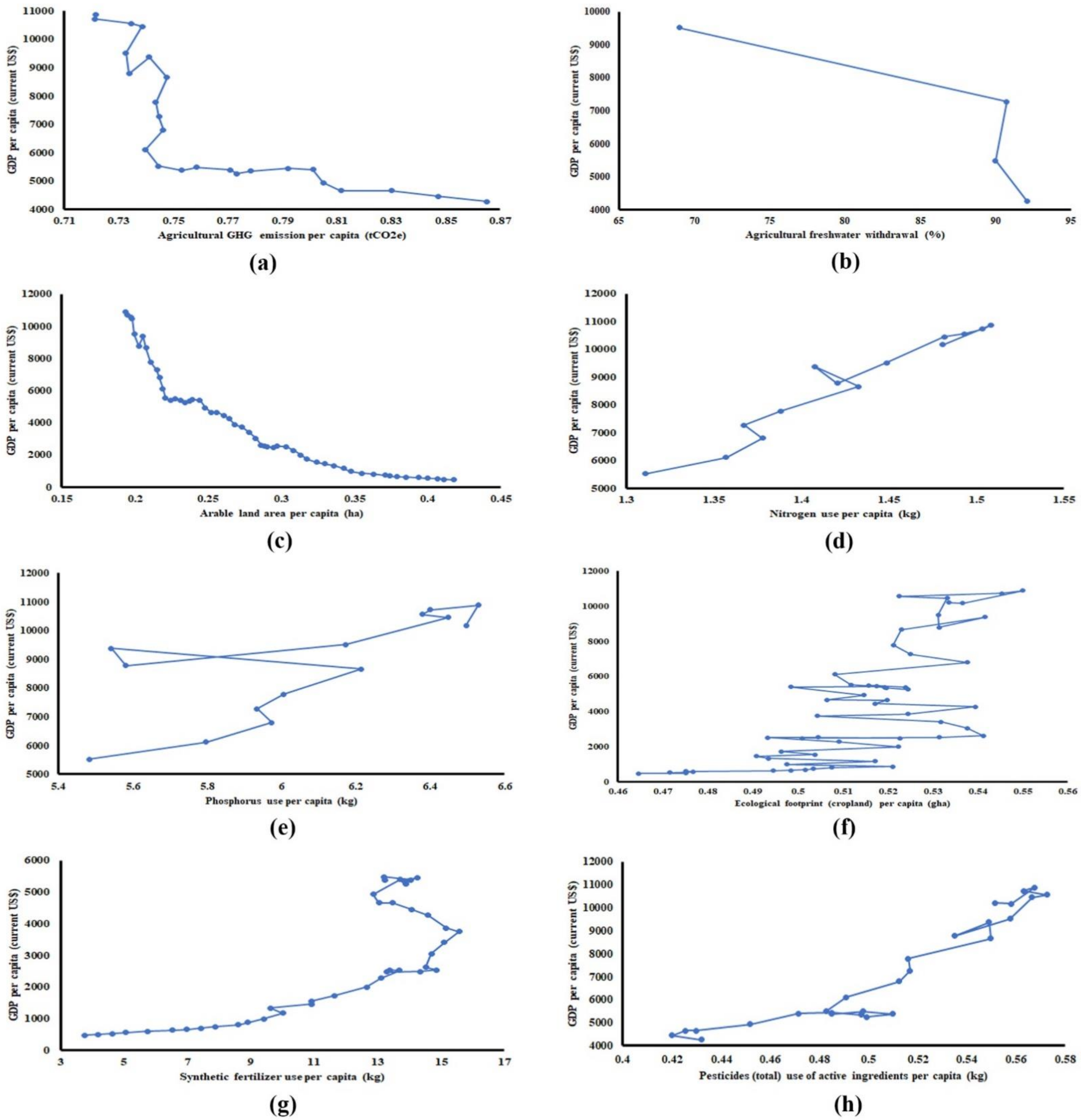


**Fig 12: Trends of change of GDP (per capita) with different indicators of planetary boundary of agriculture in the World.**

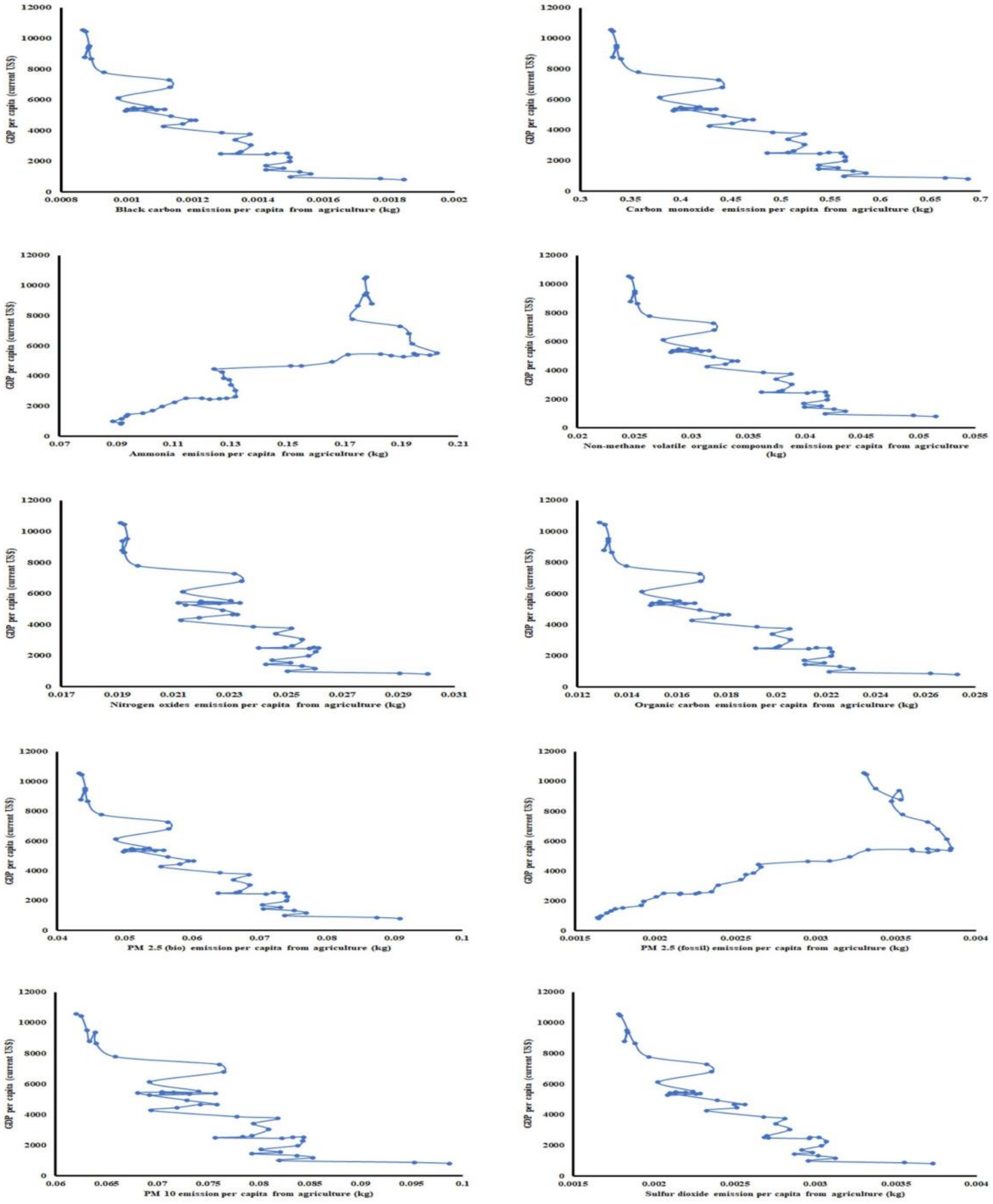


**Fig 13: Trends of change of GDP (per capita) with indicators of atmospheric pollutants by agriculture in the World.**

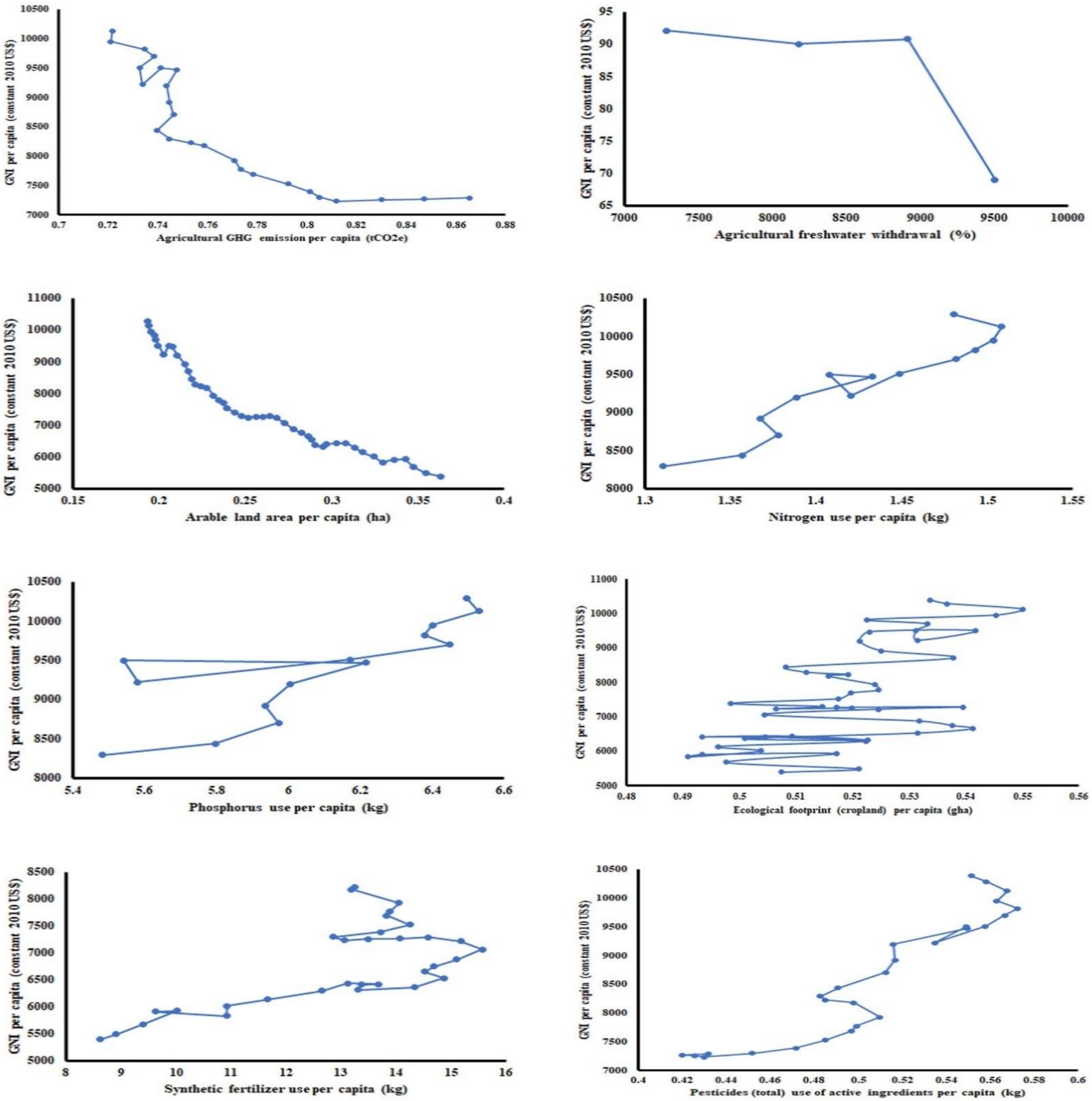


**Fig 14: Trends of change of GNI (per capita) with different indicators of planetary boundary of agriculture in the World.**

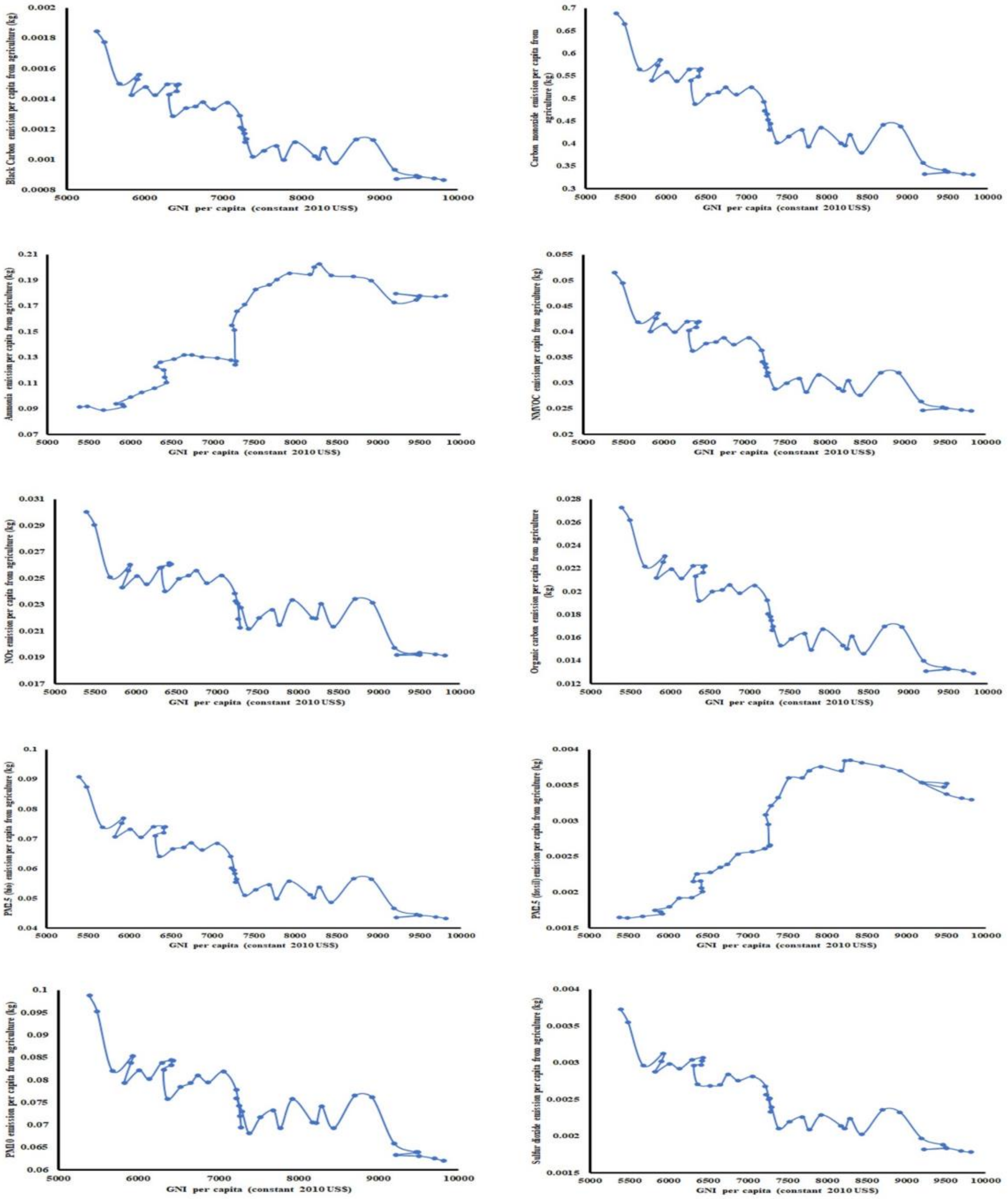


**Fig 15: Trends of change of GNI per capita (constant US$) with indicators of atmospheric pollutants by agriculture in the World.**

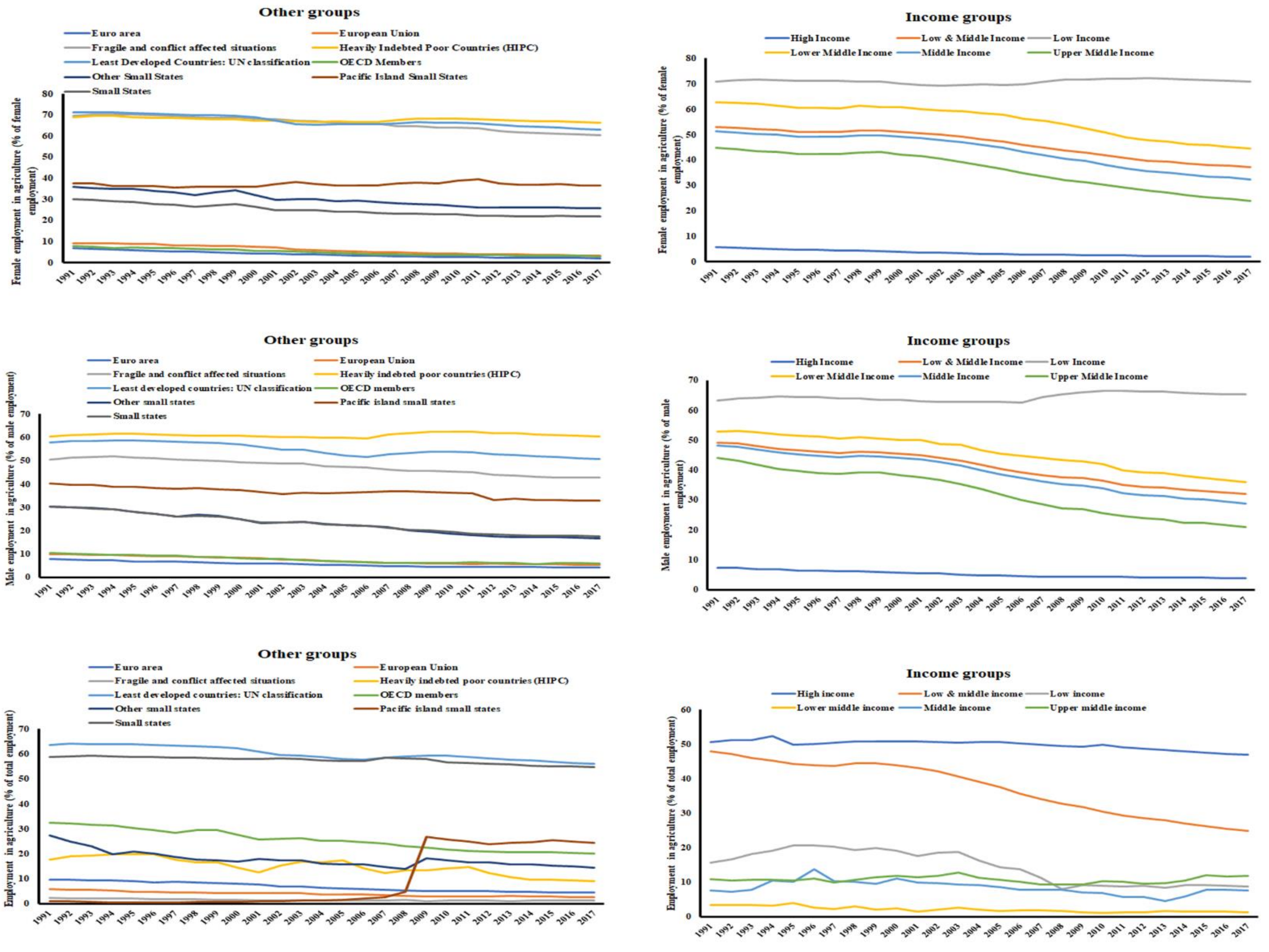


**Fig 16: Trends of socioeconomic development indicators related to agriculture in the World.**

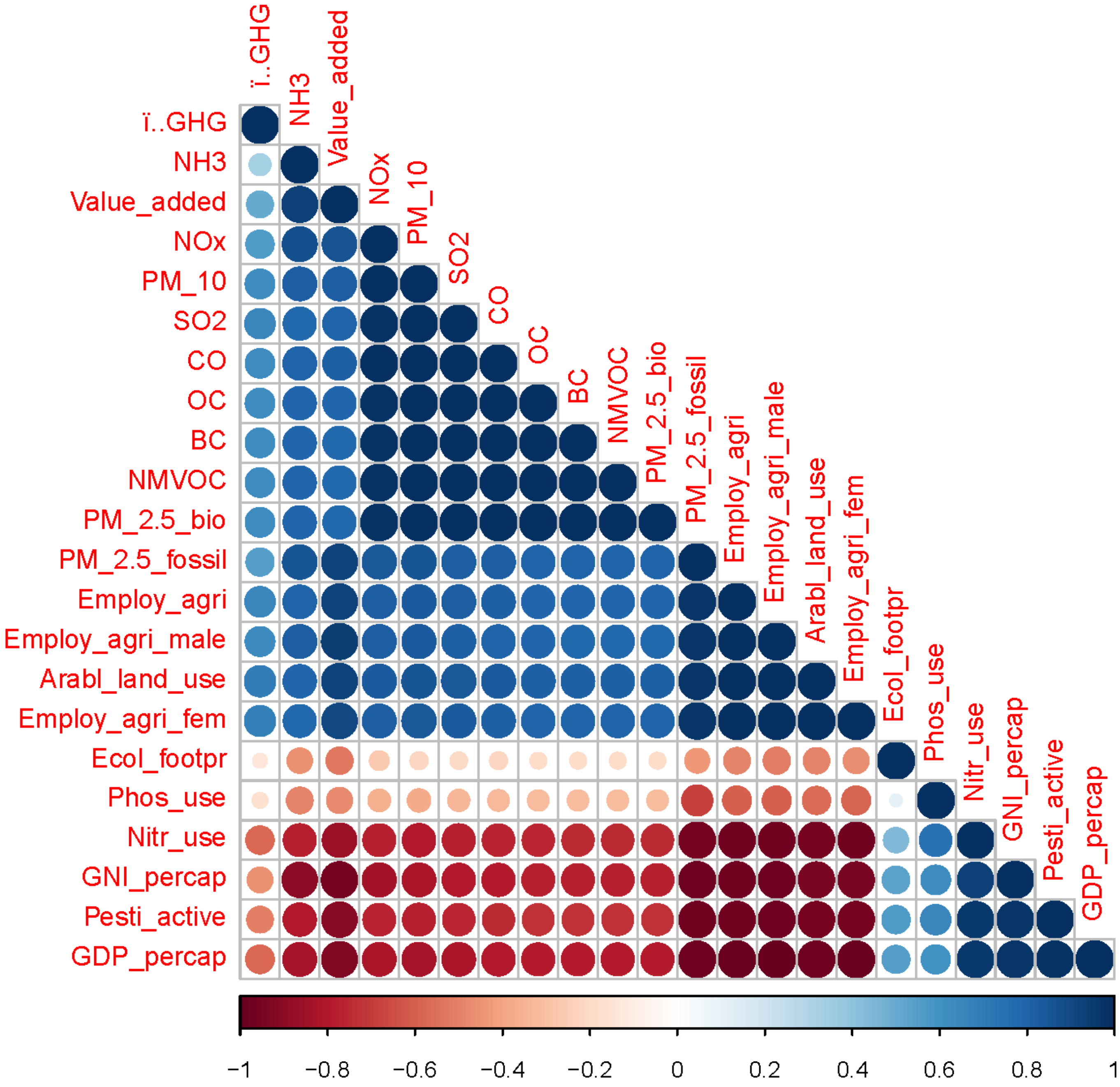


**Fig 17: Correlogram of 22 indicators related to agriculture in the World. It has been done using open source software 'R'.**
